# Firing rate adaptation affords place cell theta sweeps, phase precession and procession

**DOI:** 10.1101/2022.11.14.516400

**Authors:** Tianhao Chu, Zilong Ji, Junfeng Zuo, Yuanyuan Mi, Wen-hao Zhang, Tiejun Huang, Daniel Bush, Neil Burgess, Si Wu

**Affiliations:** School of Psychological and Cognitive Sciences, IDG/McGovern Institute for Brain Research, Center of Quantitative Biology, Peking-Tsinghua Center for Life Sciences, Academy for Advanced Interdisciplinary Studies, Peking University, China; Institute of Cognitive Neuroscience, University College London, UK; Department of Psychology Tsinghua University, China; Lyda Hill Department of Bioinformatics, O’Donnell Brain Institute, UT Southwestern Medical Center, USA; School of Computer Science, Peking University, China; Department of Neuroscience, Physiology and Pharmacology, University of College London, UK

## Abstract

Hippocampal place cells in freely moving rodents display both theta phase precession and procession, which is thought to play important roles in cognition, but the neural mechanism for producing theta phase shift remains largely unknown. Here we show that firing rate adaptation within a continuous attractor neural network causes the neural activity bump to oscillate around the external input, resembling theta sweeps of decoded position during locomotion. These forward and backward sweeps naturally account for theta phase precession and procession of individual neurons, respectively. By tuning the adaptation strength, our model explains the difference between “bimodal cells” showing interleaved phase precession and procession, and “unimodal cells” in which phase precession predominates. Our model also explains the constant cycling of theta sweeps along different arms in a T-maze environment, the speed modulation of place cells’ firing frequency, and the continued phase shift after transient silencing of the hippocampus. We hope that this study will aid an understanding of the neural mechanism supporting theta phase coding in the brain.

## Introduction

One of the strongest candidates for temporal coding of a cognitive variable by neural firing is the ‘theta phase precession’ shown by hippocampal place cells. As an animal runs through the firing field of a place cell, the cell fires at progressively earlier phases in successive cycles of the ongoing LFP theta oscillation, so that firing phase correlates with distance traveled (***O’Keefe and Recce, 1993***; ***Skaggs et al., 1996***) (see also (***Schmidt et al., 2009***)) (Fig. 1a&b). At the population level, phase precession of individual cells gives rise to forward theta sequences once starting phases are aligned across the population (***Feng et al., 2015***), where neurons representing successive locations along the trajectory of the animal display predictable firing sequences within individual theta cycles (***Johnson and Redish, 2007***). These prospective sequential experiences (looking into the future) are potentially useful for a range of cognitive faculties, e.g., planning, imagination, and decision-making (***O’Keefe and Recce, 1993***; ***Skaggs et al., 1996***; ***Hassabis et al., 2007***; ***Wikenheiser and Redish, 2015***; ***Kay et al., 2020***).

**Figure 1.**
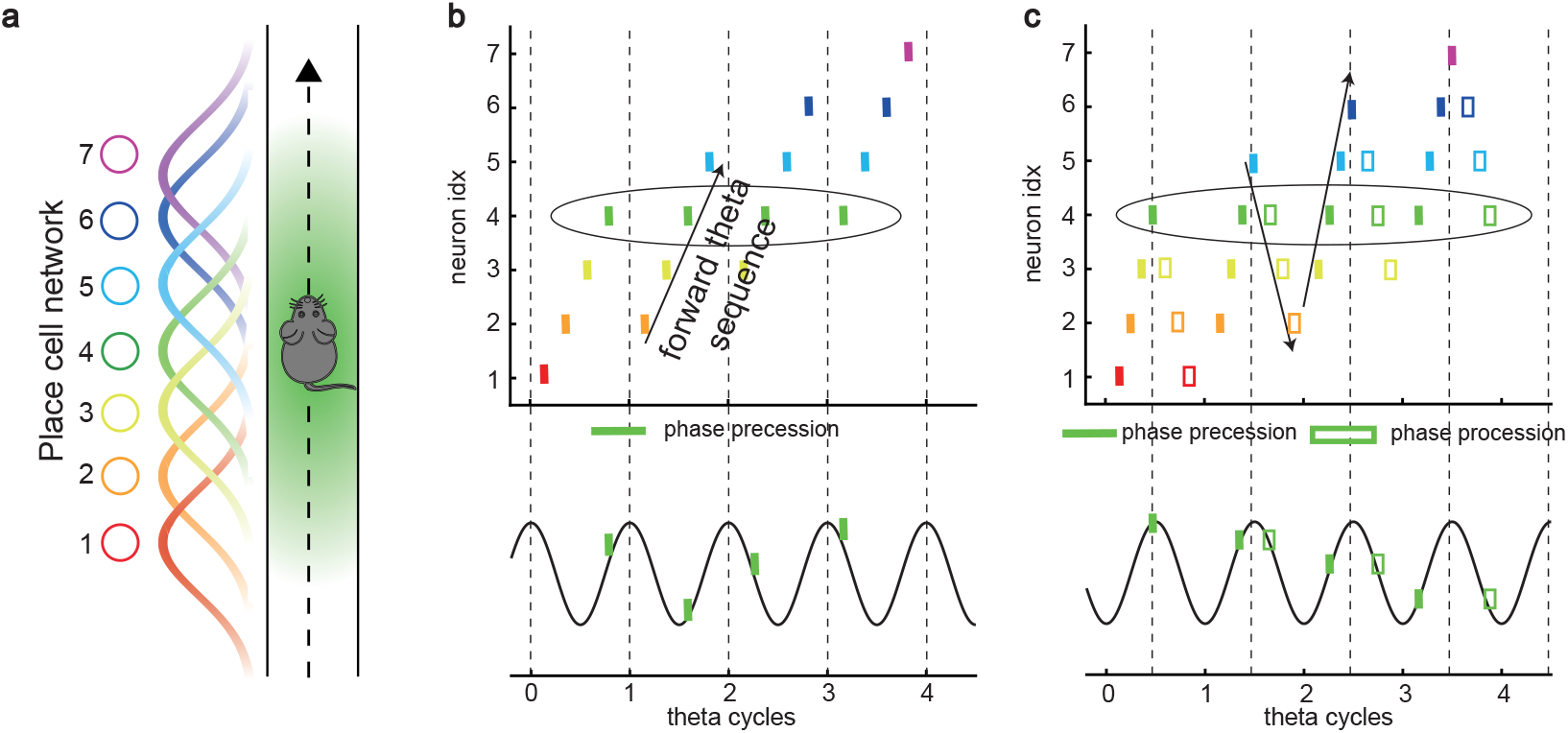
Theta sequence and theta phase shift of place cell firing. **a**, An illustration of an animal running on a linear track. A group of place cells each represented by a different color are aligned according to their firing fields on the linear track. **b**, An illustration of the forward theta sequences of the neuron population (upper panel), and the theta phase precession of the 4th place cell (represented by the green color, lower panel). **c**, An illustration of both forward and reverse theta sequences (upper panel), and the corresponding theta phase precession and procession of the 4th place cell (lower panel). The sinusoidal trace illustrates the theta rhythm of local field potential (LFP), with individual theta cycles separated by vertical dashed lines.

Besides prospective representation, flexible behaviors also require retrospective representation of sequential experiences (looking into the past). For instance, in goal-directed behaviors, it is important to relate the reward information that might only occur at the end of a sequence of events to preceding events in the sequence (***Foster et al., 2000***; ***Foster and Wilson, 2006***; ***Diba and Buzsáki, 2007***). A recent experimental study (***Wang et al., 2020***) described retrospective sequences during online behaviors (also indicated by (***Skaggs et al., 1996***; ***Yamaguchi et al., 2002***)), namely, reverse theta sequences, interleaved with forward theta sequences in individual theta cycles (Fig. 1c). Such retrospective sequences, together with the prospective sequences, may cooperate to establish higher-order associations in episodic memory (***Diba and Buzsáki, 2007***; ***Jaramillo and Kempter, 2017***; ***Pfeiffer, 2020***).

While a large number of computational models of phase precession and the associated forward theta sequences have been proposed, e.g., the single cell oscillatory models (***O’Keefe and Recce, 1993***; ***Kamondi et al., 1998***; ***Harris et al., 2002***; ***Lengyel et al., 2003***; ***Losonczy et al., 2010***) and recurrent activity spreading models (***Tsodyks et al., 1996***; ***Romani and Tsodyks, 2015***), the underlying neural mechanism for interleaved forward- and reverse-ordered sequences remains largely unclear. Do reverse theta sequences share the same underlying neural mechanism as forward sequences, or do they reflect different mechanisms? If they do, what kind of neural architecture can support the emergence of both kinds of theta phase shift? Furthermore, since forward theta sequences are commonly seen, but reverse theta sequences are only seen in some circumstances (***Wang et al., 2020***), are they commensurate with forward theta sequences? If not, to what degree are forward theta sequences more significant than the reverse ones?

To address these questions, we built a continuous attractor neural network (CANN) of the hippocampal place cell population (***Amari, 1977***; ***Tsodyks and Sejnowski, 1995***; ***Samsonovich and Mc-Naughton, 1997***; ***Tsodyks, 1999***). The CANN conveys a map of the environment in its recurrent connections that affords a single bump of activity on a topographically organized sheet of cells which can move smoothly so as to represent the location of the animal as it moves in the environment. Each neuron exhibits firing rate adaptation which destabilizes the bump attractor state. When the adaptation is strong enough, the network bump can travel spontaneously in the attractor space, which we term as the intrinsic mobility. Intriguingly, we show that, under competition between the intrinsic mobility and the extrinsic mobility caused by location-dependent sensory inputs, the network displays an oscillatory tracking state, in which the network bump sweeps back and forth around the external sensory input. This phenomenon naturally explains the theta sweeps found in the hippocampus (***Skaggs et al., 1996***; ***Burgess et al., 1994***; ***Foster and Wilson, 2007***), where the decoded position sweeps around the animal’s physical position at theta frequency. More specifically, phase precession occurs when the bump propagates forward while phase procession occurs when the network bump propagates backward. Moreover, we find that neurons can exhibit either only predominant phase precession (unimodal cells) when adaptation is relatively strong, or interleaved phase precession and procession (bimodal cells) when adaptation is relatively weak.

In addition to theta phase shift, our model also successfully explains the constant cycling of theta sweeps along different upcoming arms in a T-maze environment (***Kay et al., 2020***), and other phenomena related to phase precession of place cells (***Geisler et al., 2007***; ***Zugaro et al., 2005***). We hope that this study facilitates our understanding of the neural mechanism underlying the rich dynamics of hippocampal neurons and lays the foundation for unveiling their computational functions.

## Results

### A network model of hippocampal place cells

To study the phase shift of hippocampal place cells, we focus on a one-dimensional (1D) continuous attractor neural network (CANN) (mimicking the animal moving on a linear track, see Fig. 2a), but generalization to the 2D case (mimicking the animal moving in a 2D arena) is straightforward (see Discussion for more details). Neurons in the 1D CANN can be viewed as place cells rearranged according to the locations of their firing fields on the linear track (measured during free exploration). The dynamics of the 1D CANN is written as

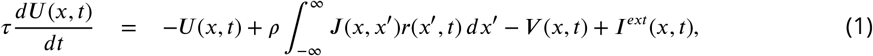

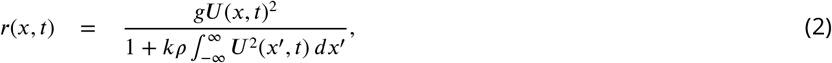

**Figure 2.**
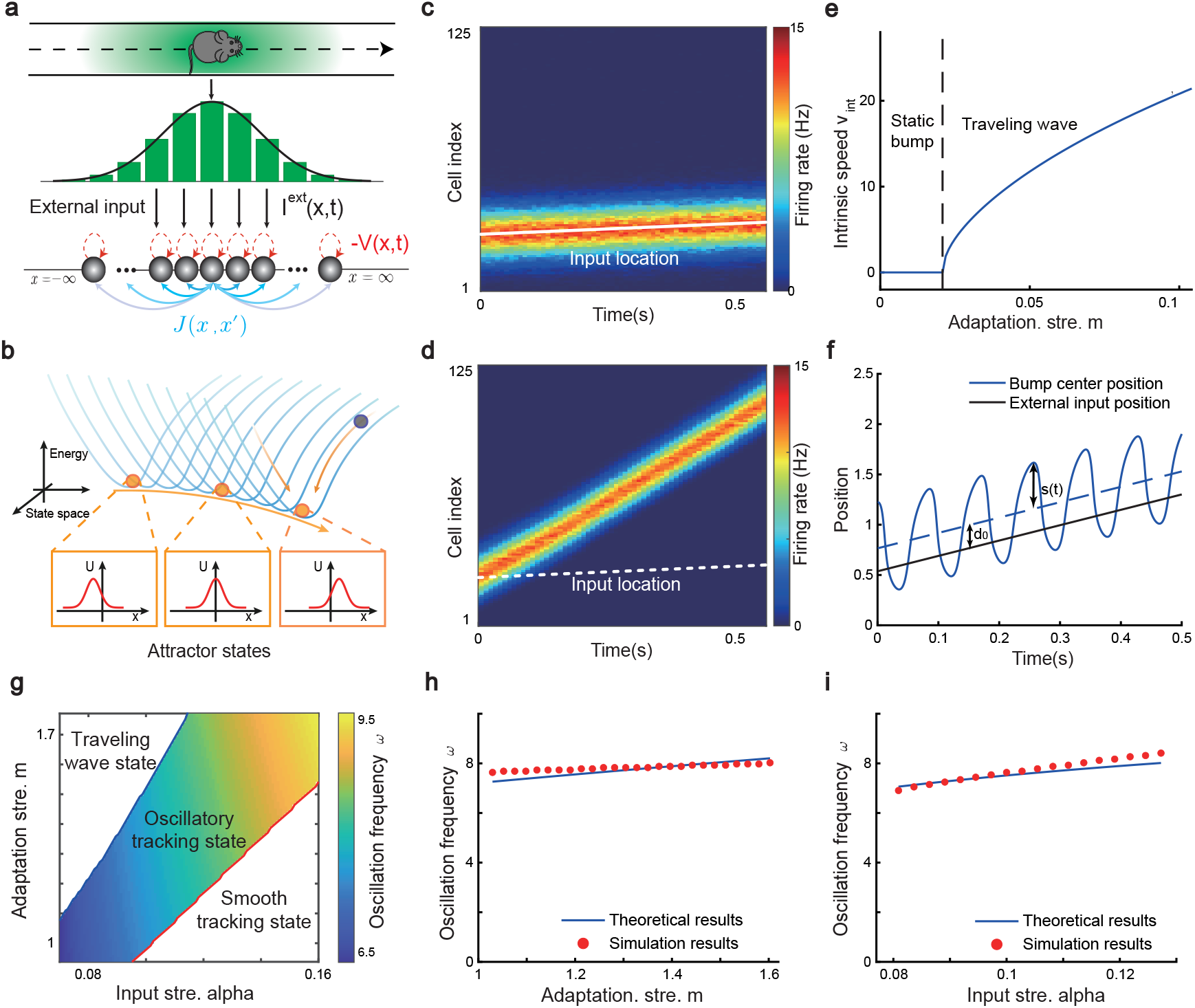
The network architecture and tracking dynamics. **a**, A 1D continuous attractor neural network formed by place cells. Neurons are aligned according to the locations of their firing fields on the linear track. The recurrent connection strength *J* (*x, x*^′^) (blue arrows) between two neurons decays with their distance on the linear track. Each neuron receives an adaptation current *−V (x, t*) (red dashed arrows). The external input *I*^*ext*^*(x, t*), represented by a Gaussian-shaped bump, conveys location-dependent sensory inputs to the network. **b**, An illustration of the state space of the CANN. The CANN holds a family of bump attractors which form a continuous valley in the energy space. **c**, The smooth tracking state. The network bump (hot colors) smoothly tracks the external moving input (the white line). The red (blue) color represents high (low) firing rate. **d**, The travelling wave state when the CANN has strong firing rate adaptation. The network bump moves spontaneously with a speed much faster than the external moving input. **e**, The intrinsic speed of the travelling wave versus the adaptation strength. **f**, The oscillatory tracking state. The bump position sweeps around the external input (black line) with an offset *d*_0_. **g**, The phase diagram of the tracking dynamics with respect to the adaptation strength *m* and the external input strength *α*. The colored area shows the parameter regime for the oscillatory tracking state. Yellow (blue) color represents fast (slow) oscillation frequency. **h-i**, Simulated (red points) and theoretical (blue line) oscillation frequency as a function of the adaptation strength (**h**) or the external input strength (**i**).

Here *U (x, t*) represents the presynaptic input to the neuron located at position *x on*the linear track, and *r*(*x, t)* represents the corresponding firing rate constrained by global inhibition (***Hao et al., 2009***). *τ* is the time constant, *ρ* the neuron density, *k* the global inhibition strength, and *g* is the gain factor. The dynamics of *U (x, t*) is determined by the leaky term *−U(x, t*), the recurrent input from other neurons, the firing rate adaptation *−V (x, t*), and the external input *I*^*ext*^*(x, t*). The recurrent connection strength *J* (*x, x*^′^) between two neurons decays with their distance. For simplicity, we set *J* (*x, x*^′^) to be the Gaussian form, i.e., 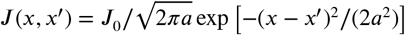, with *J*_0_ controlling the connection strength and *a* the range of neuronal interaction. Such connectivity gives rise to a synaptic weight matrix with the property of translation invariance. Together with the global inhibition, the translation invariant weight matrix ensures that the network can hold a continuous family of stationary states (attractors) when no external input and adaptation exist (***Tsodyks and Sejnowski, 1995***; ***Samsonovich and McNaughton, 1997***; ***McNaughton et al., 2006***; ***Wu et al., 2008***), where each attractor is a localized firing bump representing a single spatial location (Fig. 2b). These bump states are expressed as (see Methods. for the parameter settings and SI.2 for the detailed mathematical derivation):

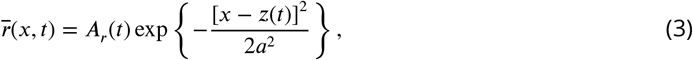

where *A*_*r*_(*t*) denotes the bump height and *z*(*t*) the bump center, i.e., the spatial location represented by the network. For convenience, we set the external input to be of the Gaussian form, which is written as *I*^*ext*^*(x, t*) = *α* exp [*−(x − v*_*ext*_*t*)^2^*/*(4*a*^2^) ], with *v*_*ext*_ representing the moving speed and *α* controlling the external input strength. Such external moving input represents location-dependent sensory inputs (i.e., corresponding to the animal’s physical location) which might be conveyed via the entorhinal-hippocampal or subcortical pathways (***Van Strien et al., 2009***). The term *−V (x, t*) represents the firing rate adaptation (***Alonso and Klink, 1993***; ***Fuhrmann et al., 2002***; ***Benda and Herz, 2003***; ***Treves, 2004***), whose dynamics is written as

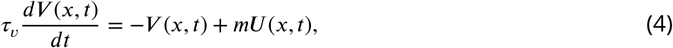

where *m* controls the adaptation strength, and *τ*_*v*_ is the time constant. The condition *τ*_*v*_ ≫ *τ* holds, implying that the firing rate adaptation is a much slower process compared to neuronal firing. In effect, the firing rate adaptation increases with the neuronal activity and contributes to destabilizing the active bump state, which induce rich dynamics of the network (see below).

### Oscillatory tracking of the network

Overall, the bump motion in the network is determined by two competing factors, i.e., the external input and the adaptation. The interplay between these two factors leads to the network exhibiting oscillatory tracking in an appropriate parameter regime. To elucidate the underlying mechanism clearly, we explore the effects of the external input and the adaptation on bump motion separately. First, when firing rate adaptation does not exist in the network (*m* = 0), the bump tracks the external moving input smoothly (see Fig. 2c). We refer to this as the “**smooth tracking state**”, where the internal location represented in the hippocampus (the bump position) is continuously tracking the animal’s physical location (the external input location). This smooth tracking property of CANNs has been widely used to model spatial navigation in the hippocampus (***Tsodyks and Sejnowski, 1995***; ***Samsonovich and McNaughton, 1997***; ***McNaughton et al., 2006***; ***Battaglia and Treves, 1998***). Second, when the external drive does not exist in the network (*α* = 0) and the adaptation strength *m* exceeds a threshold (*m* > *τ/τ*_*v*_), the bump moves spontaneously with a speed calculated as 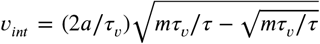 (see Fig. 2d&e and Methods. for more details). We refer to this as the “**travelling wave state**”, where the internal representation of location in the hippocampus is sequentially reactivated without external drive, resembling replay-like dynamics during a quiescent state (see Discussion for more details). This intrinsic mobility of the bump dynamics can be intuitively understood as follows. Neurons around the bump center have the highest firing rates and hence receive the strongest adaptation. Such strong adaptation destabilizes the bump stability at the current location, and hence pushes the bump away. After moving to a new location, the bump will be continuously pushed away by the firing rate adaptation at the new location. As a result, the bump keeps moving on the linear track. Similar mechanisms have been applied to explain mental exploration (***Hopfield, 2010***), preplay during sharp wave-ripple events in the hippocampus (***Azizi et al., 2013***), and the free memory recall phenomenon in the brain (***Dong et al., 2021***).

When both the external input and adaptation are applied to the CANN, the interplay between the extrinsic mobility (caused by the external input) and the intrinsic mobility (caused by the adaptation) will induce three different dynamical behaviors of the network (see **video 1** for demonstration), i.e., 1) when *m* is small and *α* is large, the network displays the smooth tracking state; 2) when *m* is large and *α* is small, the network displays the travelling wave state; 3) when both *m* and *α* have moderate values, the network bump displays an interesting state, called the “**oscillatory tracking state**”, where the bump tracks the external moving input in an oscillatory fashion (Fig. 2f&g). Intuitively, the mechanism for oscillatory tracking can be understood as follows. Due to the intrinsic mobility of the network, the bump tends to move at its own intrinsic speed (which is faster than the external moving input, see Fig. 2d), i.e., the bump tries to escape from the external input. However, due to the strong locking effect of the external input, the bump can not run too far away from the location input, but instead, is attracted back to the location input. Once the bump returns, it will keep moving in the opposite direction of the external input until it is pulled back by the external input again. Over time, the bump will sweep back and forth around the external moving input, displaying the oscillatory tracking behavior. It is noteworthy that the activity bump does not live within a window circumscribed by the external input bump (bouncing off the interior walls of the input during the oscillatory tracking state), but instead is continuously pulled back and forth by the external input (see Fig. S1).

Our study shows that during oscillatory tracking, the bump shape is roughly unchanged (see previous sections for the condition of shape variability), and the bump oscillation can be well represented as the bump center sweeping around the external input location. The dynamics of the bump center can be approximated as a propagating sinusoidal wave (Fig. 2f), i.e.,

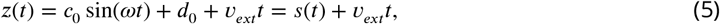

where *z*(*t*) is the bump center at time *t* (see Eq. 3). *s(t)* denotes the displacement between the bump center and the external input, which oscillates at the frequency *ω* with the amplitude *c*_0_ > 0 and a constant offset *d*_0_ > 0 (see Methods. for the values of these parameters and SI.3 for the detailed derivation). When the firing rate adaptation is relatively small, the bump oscillation frequency can be analytically solved to be (see also Fig. S2):

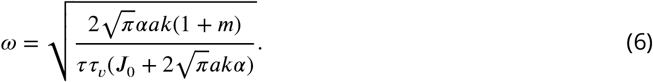

We see that the bump oscillation frequency *ω*increases sublinearly with the external input strength *α* and the adaptation strength *m* (Fig. 2h&i). By setting the parameters appropriately, the bump can oscillate in the theta band (6-10 Hz), thus approximating the experimentally observed theta sweeps (see below). Notably, LFP theta is not explicitly modelled in the network. However, since theta sweeps are bounded by individual LFP theta cycles in experiments, they share the same oscillation frequency as LFP theta. For convenience, we will frequently use the term LFP theta below and study firing phase shift in individual oscillation cycles.

### Oscillatory tracking accounts for both theta phase precession and procession of hippocampal place cells

In our model, the bump center and external input represent the decoded and physical positions of the animal, respectively, thus the oscillatory tracking of the bump around the external input naturally gives rise to the forward and backward theta sweeps observed empirically (Fig. 3a&b) (***Wang et al., 2020***). Here we show that oscillatory tracking of the bump accounts for the theta phase precession and procession of place cell firing.

**Figure 3.**
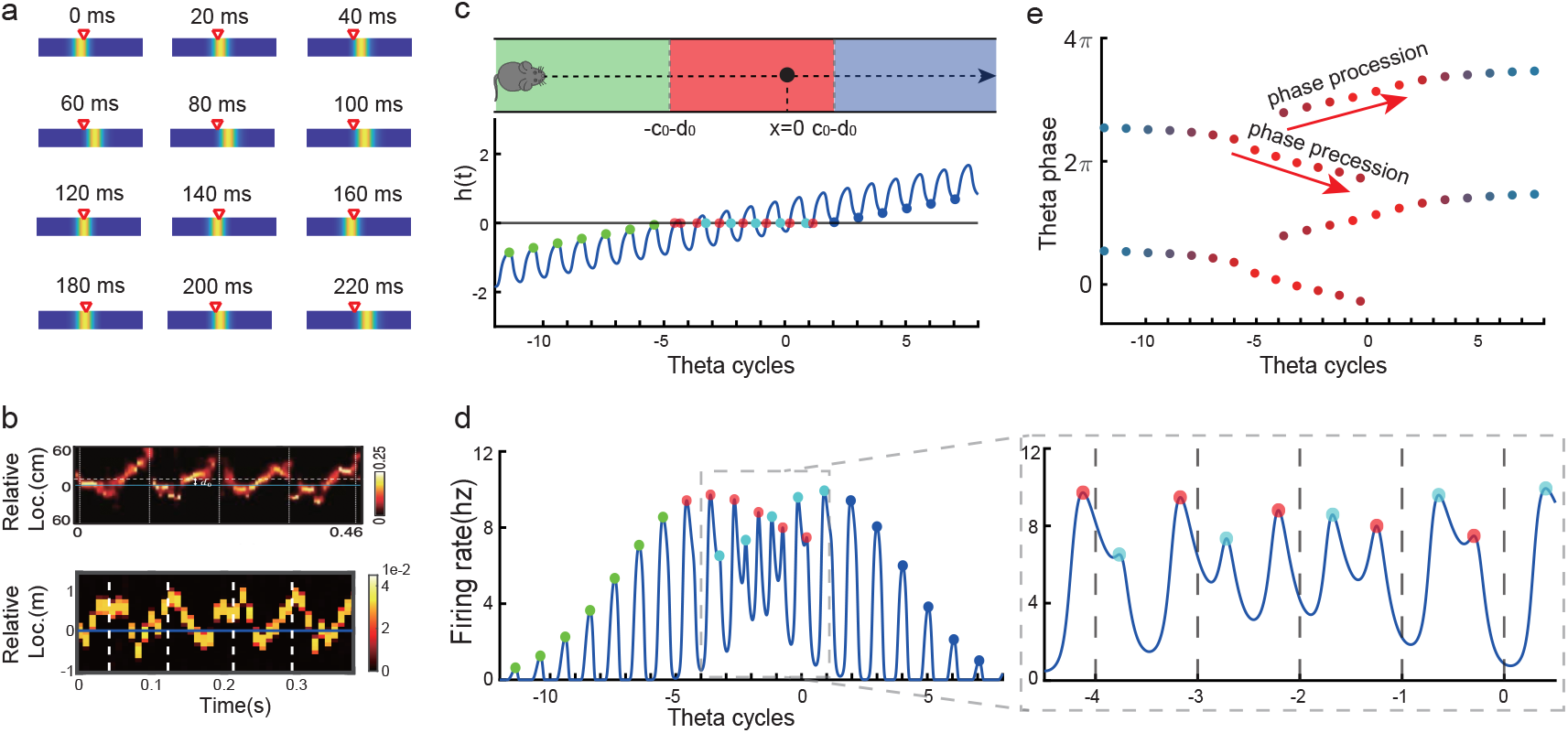
Oscillatory tracking accounts for theta sweeps and theta phase shift. **a**, Snapshots of the bump oscillation along the linear track in one theta cycle (0 ms - 140 ms). Red triangles indicate the location of the external moving input. **b**, Decoded relative positions based on place cell population activities. Upper panel: experimental data, adapted from ***Wang et al. (2020***). Lower panel: the relative locations of the bump center (shown by the 10 most active neurons at each timestamp) with respect to the location of the external input (horizontal line) in five theta cycles. **c**, Upper panel: the process of the animal running through the firing field of the probe neuron (large black dot) is divided into three stages: the entry stage (green), the phase shift stage (red) and the departure stage (blue). Lower panel: the displacement between the bump center and the probe neuron as the animal runs through the firing field. The horizontal line represents the location of the probe neuron, which is *x* = 0. **d**, The firing rates of the probe neuron as the animal runs through the firing field. Colored points indicate firing peaks. The trace of the firing rate in the phase shift stage (the dashed box) is enlarged in the sub-figure on the right hand-side, which exhibits both phase precession (red points) and procession (blue points) in successive theta cycles. **e**, The firing phase shift of the probe neuron in successive theta cycles. Red points progress to earlier phases from *π*/2 to −*π*/2 and blues points progress to later phases from *π*/2 to 3*π*/2. The color of the dots represent the peak firing rates, which is also shown in d.

Without loss of generality, we select the neuron at location *x* = 0 as the probe neuron and examine how its firing phase changes as the external input traverses its firing field (Fig. 3c). In the absence of explicitly simulated spike times, the firing phase of a neuron in each theta cycle is measured by the moment when the neuron reaches the peak firing rate (see Methods. for modeling spike times in the CANN). Based on Eqs. 3 & 5, the firing rate of the probe neuron, denoted as *r*_0_(*t*), is expressed as

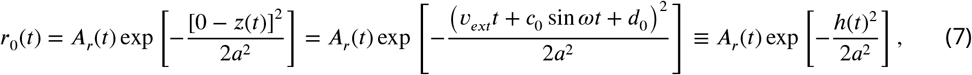

where *A*_*r*_ *(t)* is the bump height, and *h*(*t*) is an oscillatory moving term denoting the displacement between the bump center and the location of the probe neuron. It is composed of a moving signal *v*_*ext*_*t*and an oscillatory signal *c*_0_ sin *ωt*+ *d*_0_, with *c*_0_ the oscillation amplitude, *ω*the frequency and *d*_0_ an oscillation offset constant. It can be seen that the firing rate of the probe neuron is determined by two factors, *A*_*r*_(*t*) and *h(t)*. To simplify the analysis below, we assume that the bump height *A*_*r*_(*t*) remains unchanged during bump oscillations (for the case of time-varying bump height, see previous sections). Thus, the firing rate only depends on *h(t)*, which is further determined by two time-varying terms, the oscillation term *c*_0_ sin *ωt* and the location of the external input *v*_*ext*_*t*. The first term contributes to firing rate oscillations of the probe neuron, and the second term contributes to the envelope of neuronal oscillations exhibiting a waxing-and-waning profile over time, as the external input traverses the firing field (the absolute value |*v*_*ext*_*t*| first decreases and then increases; see Fig. 3d, also **video 2**). Such a waxing-and-waning profile agrees well with the experimental data (***Skaggs et al., 1996***). In each LFP theta cycle, the peak firing rate of the probe neuron is achieved when |*h(t)*| reaches a local minima (Fig. 3c&d). We differentiate three stages as the external input passes through the probe neuron (i.e., the animal travels through the place field of the probe neuron), i.e.,

- **the entry stage.** As the external input enters the firing field of the probe neuron (moving from left to right), *h(t)* < 0 always holds (Fig. 3c). In this case, the peak firing rate of the probe neuron in each oscillatory cycle is achieved when *h(t)* reaches the maximum (i.e., |*h(t)*| reaches the minimum). This corresponds to *c*_0_ sin *ωt = c*_0_, i.e., *ωt = π/*2 (Fig. 3e). This means that the firing phase of the probe neuron at the entry stage is constant, which agrees with experimental observations (***O’Keefe and Recce, 1993***; ***Skaggs et al., 1996***).
- **the phase shift stage.** As the external input moves into the centre of the firing field, *h(t) =* 0 can be achieved in each oscillatory cycle (Fig. 3c). Notably, it is achieved twice in each cycle, once as the bump sweeps over the probe neuron in the forward direction and the other as the bump sweeps over the probe neuron in the backward direction. Therefore, there are two firing peaks in each bump oscillation cycle (Fig. 3d), which are expressed as (by solving *v*_*ext*_*t* + *c*_0_ sin*ωt*+ *d*_0_ = 0):

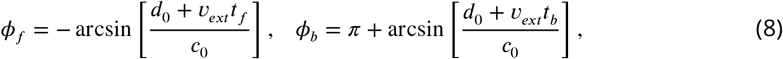

where *t*_*f*_ and *t*_*b*_ denote the moments of peak firing in the forward and backward sweeps, respectively, and *ϕ*_*f*_ and *ϕ*_*b*_ the corresponding firing phases of the probe neuron. As the external input travels from (*−c*_0_ *− d*_0_) to (*c*_0_ *− d*_0_), the firing phase *ϕ*_*f*_ in the forward sweep decreases from *π/*2 to −*π*/2, while the firing phase *ϕ*_*r*_ *in* the backward sweep increases from *π/*2 to 3*π*/2 (Fig. 3e). These give rise to the phase precession and procession phenomena, respectively, agreeing well with experimental observations (***Skaggs et al., 1996***; ***Wang et al., 2020***; ***Yamaguchi et al., 2002***).
- **the departure stage.** As the external input leaves the firing field, *h(t)* > 0 always holds (Fig. 3c), and the peak firing rate of the probe neuron is achieved when *h(t)* reaches its minimum in each oscillatory cycle, i.e., *c*_0_ sin(*ωt) = −c*_0_ with *ωt = π/*2 (Fig.3e). Therefore, the firing phase of the probe neuron is also constant during the departure stage

In summary, oscillatory tracking of the CANN well explains the firing phase shift of place cells when the animal traverses their firing fields. Specifically, when the animal enters the place field, the firing phase of the neuron remains constant, i.e., no phase shift occurs, which agrees with experimental observations (***O’Keefe and Recce, 1993***; ***Skaggs et al., 1996***). As the animal approaches the centre of the place field, the firing phase of the neuron starts to shift in two streams, one to earlier phases during the forward sweeps and the other to later phases during the backward sweeps. Finally, when the animal leaves the place field, the firing phase of the neuron stops shifting and remains constant. Over the whole process, the firing phase of a place cell is shifted by 180 degrees, which agrees with experimental observations (***O’Keefe and Recce, 1993***; ***Skaggs et al., 1996***).

### Different adaptation strengths account for bimodal and unimodal cells

The results above show that during oscillatory tracking, a place cell exhibits both significant phase precession and procession, which are associated with two firing peaks in a theta cycle. These neurons have been described as bimodal cells (***Wang et al., 2020***) (Fig. 4a). Conversely, previous experiments have primarily focused on the phase precession of place cell firing, while tending to ignore phase procession, which is a relatively weaker phenomenon (***O’Keefe and Recce, 1993***; ***Skaggs et al., 1996***). Place cells with negligible phase procession have been described as unimodal cells (Fig. 4b).

**Figure 4.**
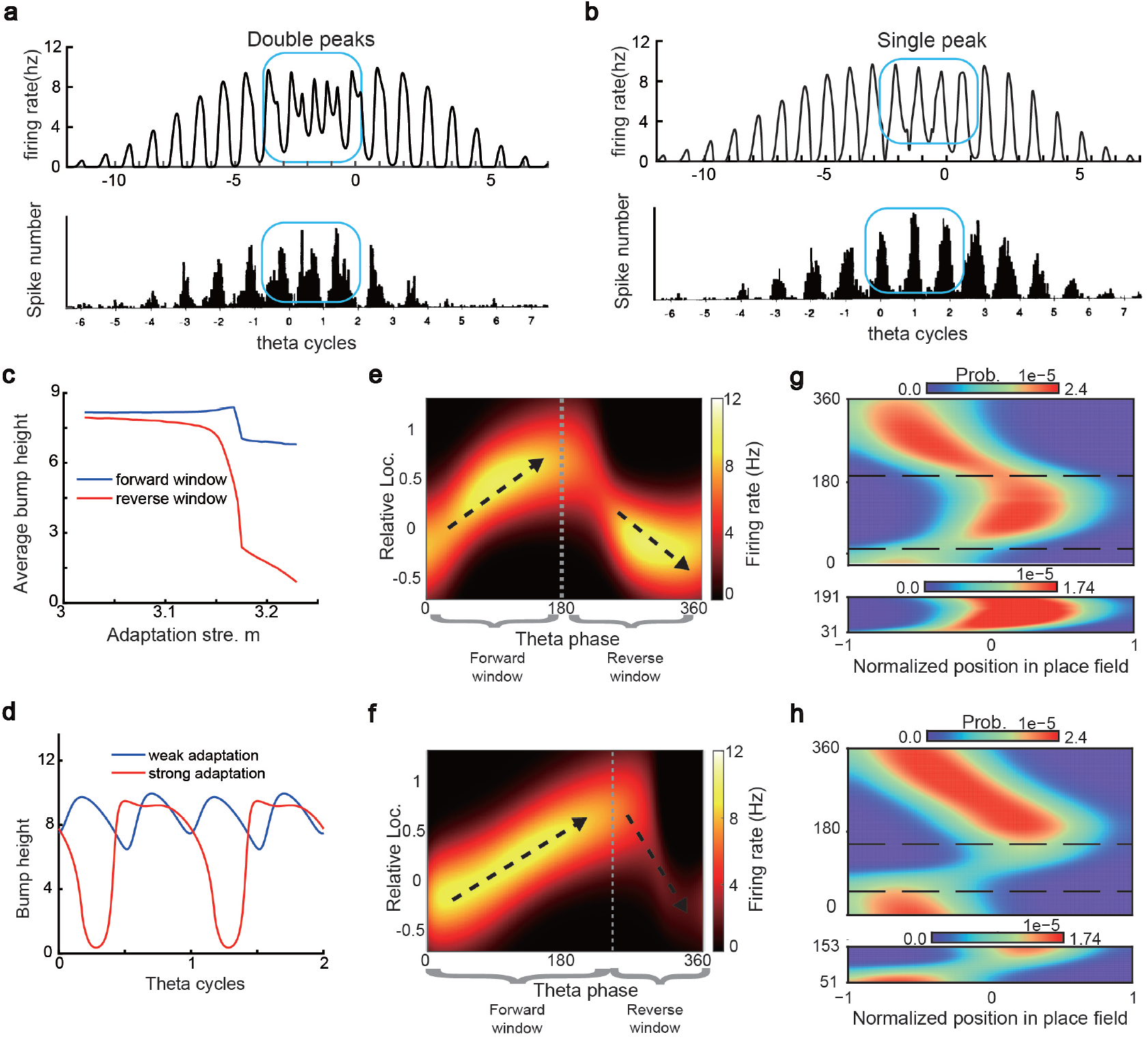
Different adaptation strengths account for the emergence of bimodal and unimodal cells. **a**, The firing rate trace of a typical bimodal cell in our model (upper panel) and the experiment data (lower panel, adapted from (***Skaggs et al., 1996***)). Blue boxes mark the phase shift stage. Note that there are two peaks in each theta cycle. **b**, The firing rate trace of a typical unimodal cell. Note that there is only one firing peak in each theta cycle. **c**, The averaged bump heights in the forward (blue curve) and backward windows (red curve) as a function of the adaptation strength *m*. **d**, Variation of the bump height when the adaptation strength is relatively small (blue line) or large (red line). **e-f,** Relative location of the bump center in a theta cycle when adaptation strength is relatively small (**e**) or large (**f**). Dashed line separate the forward and backward windows. **g-h**, Theta phase as a function of the normalized position of the animal in place field, averaged over all bimodal cells (**g**) or over all unimodal cells (**h**). *−*1 indicates that the animal just enters the place field, and 1 represents that the animal is about to leave the place field. Dashed lines separate the forward and backward windows. The lower panels in both **g** and **h** present the rescaled colormaps only in the backward window.

Here, we show that by adjusting a single parameter in the model, i.e., the adaptation strength *m*, neurons in the CANN can exhibit either interleaved phase precession and procession (bimodal cells) or predominant phase precession (unimodal cells). To understand this, we first recall that the firing rate adaptation is a much slower process compared to neural firing and its timescale is in the same order as the LFP theta (i.e., *τ*_*v*_ = 100 ms while *τ* = 5 ms). This implies that when the bump sweeps over a neuron, the delayed adaptation it generates will suppress the bump height as it sweeps back to the same location. Furthermore, since the oscillatory tracking always begins with a forward sweep (as the initial sweep is triggered by the external input moving in the same direction), the suppression effects are asymmetric, that is, forward sweeps always strongly suppress backward sweeps. On the contrary, the opposite effect is much smaller, since neuronal activities in backward sweeps have already been suppressed, and they can only generate weak adaptation. Because of this asymmetric suppression, the bump height in the forward sweep is always higher than that in the backward sweep (see Fig. 4c and Fig. S3a ). When the adaptation strength *m* is small, the suppression effect is not significant, and the attenuation of the bump height during the backward sweep is small (Fig. 4d). In such case, the firing behavior of a place cell is similar to the situation as the bump height remains unchanged as analyzed in previous sections, i.e., the neuron can generate two firing peaks in a theta cycle at the phase shift stage, manifesting the property of a bimodal cell of having both significant phase precession and procession (Fig. 4e&g and **video 2** ). When the adaptation strength *m* is large, the bump height in the backward sweep attenuates dramatically (see Fig. 4c&d and the video demonstration). As a result, the firing peak of a place cell in the backward sweep becomes nearly invisible at the phase shift stage, and the neuron exhibits only predominant phase precession, manifesting the property of a unimodal cell (Fig. 4f&h and **video 3**).

In summary, different adaptation strengths explain the emergence of bimodal and unimodal cells. In fact, there is no sharp separation between bimodal and unimodal cells. As the firing rate adaptation gets stronger, the network bump is more attenuated during the backwards sweep, and cells with the bimodal firing property will gradually behave more like those with the unimodal firing property (see Fig. S3b). Moreover, our model confirms that even though phase procession is weak, it still exist in unimodal cells (Fig. 4h lower panel), which has been reported in previous studies (***Wang et al., 2020***; ***Yamaguchi et al., 2002***). This implies that phase procession is not a characteristic feature of bimodal cells, but instead, is likely a common feature of hippocampal activity, with a strength controlled by adaptation. Furthermore, the experimental data (***Fernández-Ruiz et al., 2017***) has indicated that there is a laminar difference between unimodal cells and bimodal cells, with bimodal cells correlating more with the firing patterns of deep CA1 neurons and unimodal cells with the firing patterns of superficial CA1 neurons. Our model suggests that this difference may come from the different adaptation strengths in the two layers.

### Constant cycling of multiple future scenarios in a T-maze environment

We have shown that our model can reproduce the forward and backward theta sweeps of decoded position when the animal runs on a linear track. It is noteworthy that there is only a single hypothetical future scenario in the linear track environment, i.e, ahead of the animal’s position, and hence place cells firing phase can only encode future positions in one direction. However, flexible behaviors requires the animal encoding multiple hypothetical future scenarios in a quick and constant manner, e.g., during decision-making and planning in complex environments (***Johnson and Redish, 2007***; ***Wikenheiser and Redish, 2015***). One recent study (***Kay et al., 2020***) showed constant cycling of theta sweeps in a T-maze environment (Fig. 5a), that is, as the animal approaches the choice point, the decoded position from hippocampal activity propagates down one of the two arms alternatively in successive LFP theta cycles. To reproduce this phenomenon, we change the structure of the CANN from a linear track shape to a T-maze shape where the neurons are aligned according to the location of their firing fields in the T-maze environment. Neurons are connected with a strength proportional to the Euclidean distance between their firing fields on the T-maze and the parameters are set such that the network is in the oscillatory tracking state (see details in Methods. ). Mimicking the experimental protocol, we let the external input (the artificial animal) move from the end of the center arm to the choice point. At the beginning, when the external input is far away from the choice point, the network bump sweeps back and forth along the center arm, similar to the situation on the linear track. As the external input approaches the choice point, the network bump starts to sweep onto left and right arms alternatively in successive theta cycles (Fig. 5b and **video 4**; see also ***Romani and Tsodyks (2015***) for a similar model of cyclical sweeps spanning several theta cycles). The underlying mechanism is straightforward. Suppose that the bump first sweeps to the left arm from the current location, it will sweep back to the current location first due to the attraction of the external input. Then in the next round, the bump will sweep to the right arm, since the neurons on the left arm are suppressed due to adaptation. This cycling process repeats constantly between the two upcoming arms before the external input enters one of the two arms (i.e, before the decision is made). At the single cell level, this bump cycling phenomenon gives rise to the “cycle skipping” effect (***Kay et al., 2020***; ***Deshmukh et al., 2010***; ***Brandon et al., 2013***), where a neuron whose place field is on one of the two arms fires on every other LFP theta cycle before the decision is made (Fig. 5c left panel and Fig. 5d upper panel). For example, a pair of cells with firing fields on each of the two arms will fire in regular alternation on every other theta cycle (Fig. 5c right panel and Fig. 5d lower panel). These cell-level firing patterns agree well with the experimental observations (***Kay et al., 2020***).

**Figure 5.**
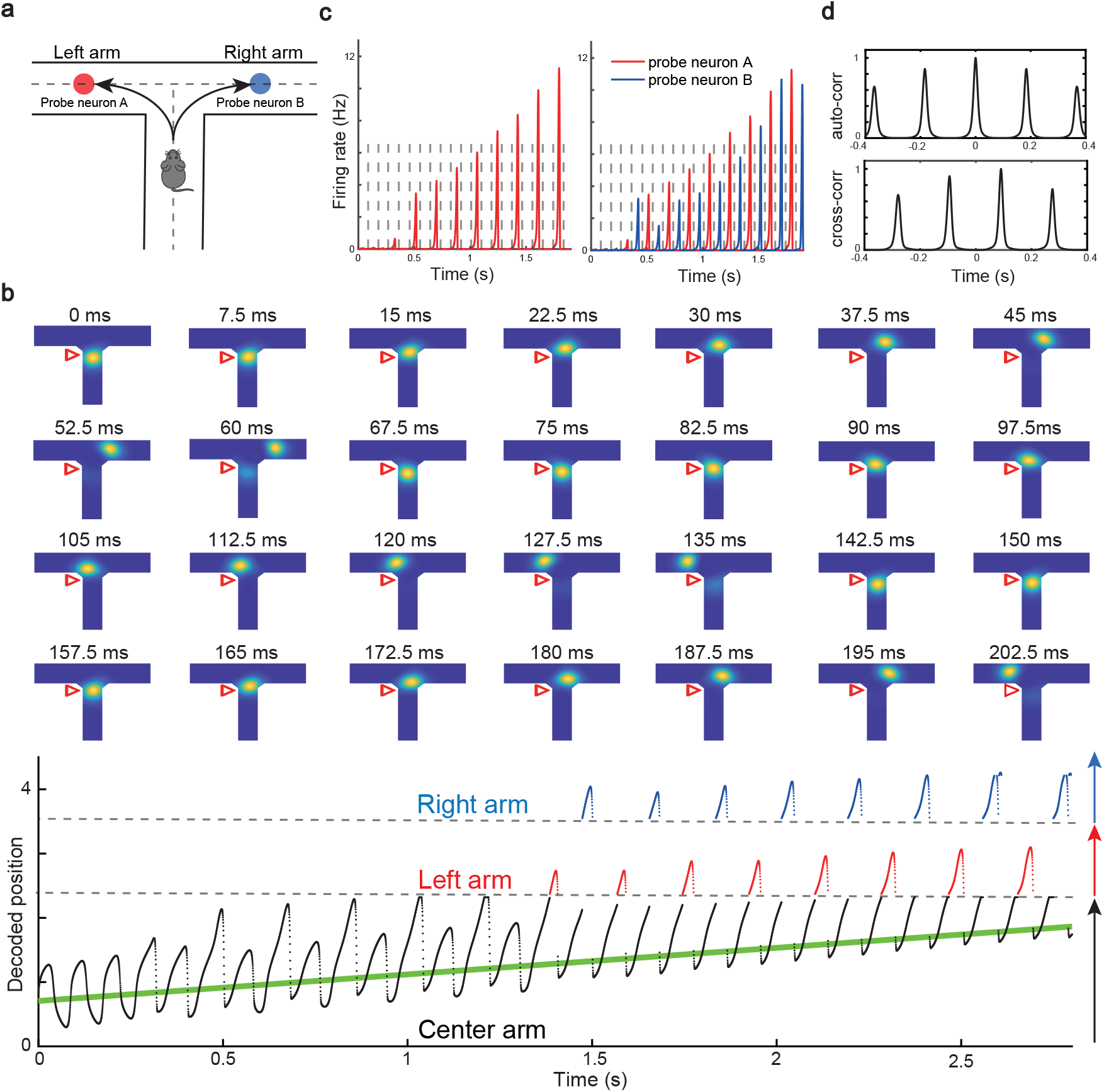
Constant cycling of future positions in a T-maze environment. **a**, An illustration of an animal navigating a T-maze environment with two possible upcoming choices (the left and right arms). **b**, Upper panel: Snapshots of constant cycling of theta sweeps on two arms when the animal is approaching the choice point. Red triangle marks the location of the external input. Note that the red triangle moves slightly towards the choice point in the 200 ms duration. Lower panel: Constant cycling of two possible future locations. The black, red and blue traces represent the bump location on the center, left and right arms, respectively. The green line marks the location of the external moving input. **c**, Left panel: the firing rate trace of a neuron A on the left arm when the animal approaches the choice point. Right panel: the firing rate traces of a pair of neurons when the animal approaches the choice point, with neuron A (red) on the left arm and neuron B (blue) on the right arm. Dashed lines separate theta cycles. **d**, Upper panel: the auto-correlogram of the firing rate trace of probe neuron A. Lower panel: the cross-correlogram between the firing rate trace of neuron A and the firing rate trace of neuron B.

In summary, our model, extended to a T-maze structure, explains the constant cycling of two possible future scenarios in a T-maze environment. The underlying mechanism relies on delayed adaptation, which alternately causes neurons on one arm to be more suppressed than those on the other arm. Such high-speed cycling may contribute to the quick and continuous sampling among multiple future scenarios in real-world decision-making and planning. We also note that there is a cyclical effect in the sweep lengths across oscillation cycles before the animal enters the left or right arm (see Fig. 5b lower panel), which may be interesting to check in the experimental data in future work (see Discussion for more details).

### Robust phase coding of position with place cells

As the firing rate shows large variability when the animal runs through the firing field (***Fenton and Muller, 1998***), it has been suggested that the theta phase shift provides an additional mechanism to improve the localization of animals (***O’keefe and Burgess, 2005***). Indeed, ***Jensen and Lisman (2000***) showed that taking phase into account leads to a significant improvement in the accuracy of localizing the animal. To demonstrate the robustness of phase coding, previous experiments showed two intriguing findings: a linear relationship between the firing frequency of place cells and the animal’s moving speed (***Geisler et al., 2007***) (Fig. 6a), and the continued phase shift after interruption of hippocampal activity (***Zugaro et al., 2005***) (Fig. 6b). We show that our model can also reproduce these two phenomena.

**Figure 6.**
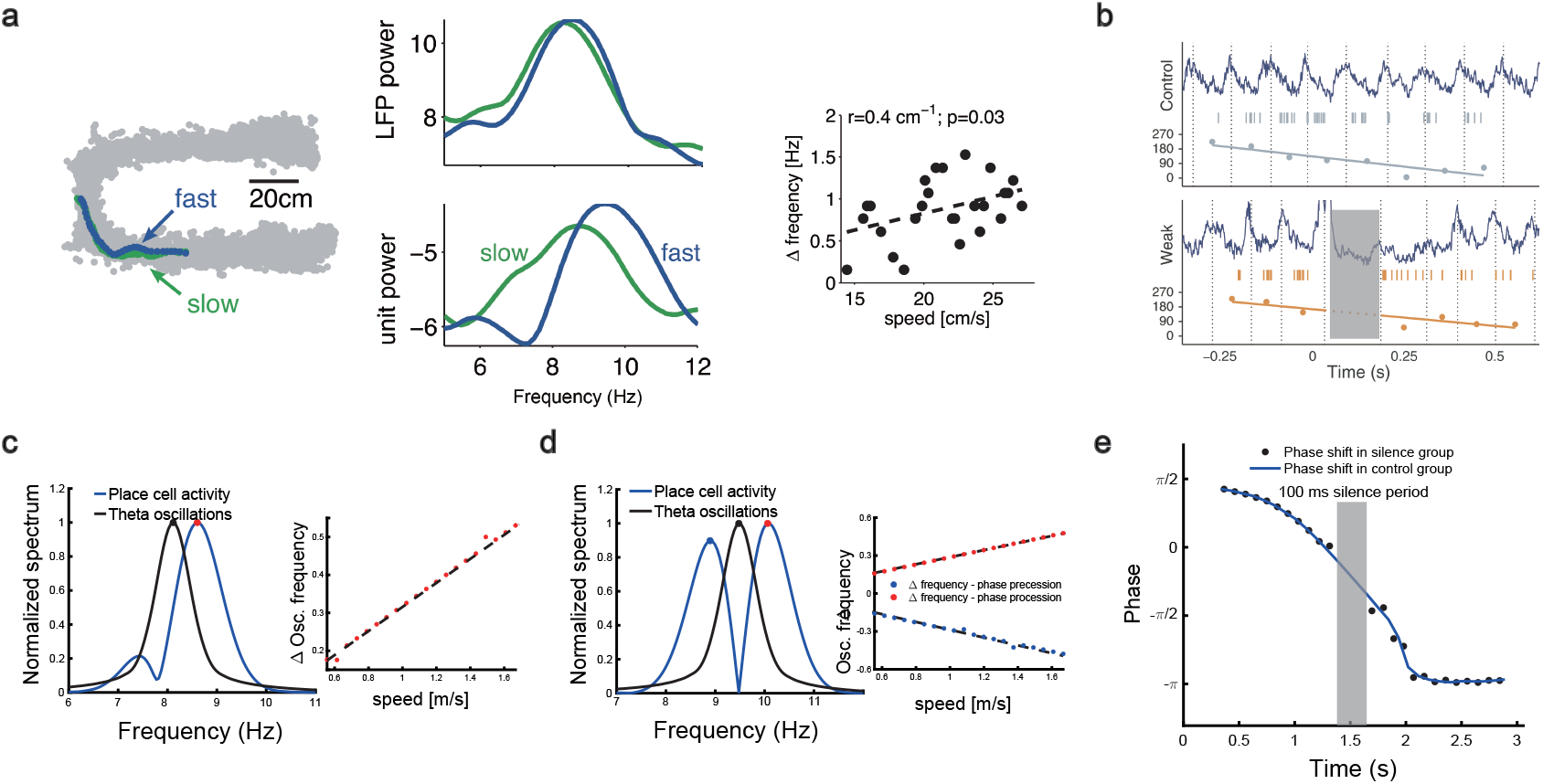
Robust phase coding of position. **a**, speed modulation of place cell firing frequency, adapted from ***Geisler et al. (2007***). Oscillation frequency of the place cell is higher in the faster run trial (blue) then that in the slower run trail (green). The frequency difference is linearly increased with the running speed. **b**, Phase precession is preserved after stimulation-induced perturbation (grey shaded area of the yellow part), adapted from ***Zugaro et al. (2005***). **c**, left: normalized spectrum of bump oscillation (black curve) and the oscillation of a unimodal cell (blue curve). Right: linear relationship between the frequency difference and the running speed. **d**, same as **c** but for a bimodal cell. **e**, silencing the network activity for 100 ms (grey shaded area) when the external moving input passes through the center part of the place field of a unimodal cell. Theta phase shifts of the unimodal cell are shown with (black points) or without (blue curve) silencing the network.

To investigate the relationship between the single cell’s oscillation frequency and the moving speed as the animal runs through the firing field, we consider a unimodal cell with predominant phase precession as studied in ***Geisler et al. (2007***). As we see from Fig. 3d and Fig. 4a&b, when the animal runs through the firing field of a place cell, the firing rate oscillates because the activity bump sweeps around the firing field center. Therefore, the firing frequency of a place cell has a baseline theta frequency, which is the same as the bump oscillation frequency. Furthermore, due to phase precession, there will be half a cycle more than the baseline theta cycles as the animal runs over the firing field, and hence single cell oscillatory frequency will be higher than the baseline theta frequency (Fig. 6c). The faster the animal runs, the faster the extra half cycle can be accomplished. Consequently, the firing frequency will increase more (a steeper slope in Fig. 6c red dots) than the baseline frequency. This linear relationship ensures that the firing phase of a unimodal cell in each theta cycle is locked with the relative location of the animal in the firing field of that cell, which supports a robust phase-position code. Notably, in our model, the speed modulation of the place cells’ firing frequency is not the cause of theta phase shift, but rather a result of oscillatory tracking. This is different from the dual oscillator model (***Lengyel et al., 2003***), which assumes that phase precession is caused by a speed-dependent increase in the dendritic oscillation frequency (see Discussion for more details).

In a different experiment, (***Zugaro et al., 2005***) found that the firing phase of a place cell continues to precess even after hippocampal activity was transiently silenced for up to 250 ms (around 2 theta cycles) (Fig. 6b). To reproduce this phenomenon, we also study a unimodal cell by manually turning off the network activity for a few hundred milliseconds (by setting *r*(*x, t)* = 0 for all neurons) and then letting the network dynamics evolves again with all parameters unchanged. Based on the theoretical analysis (Eq. 8), we see that the firing phase is determined by the location of the animal in the place field, i.e., *v*_*ext*_*t*. This means that the firing phase keeps tracking the animal’s physical location. No matter how long the network is inactivated, the new firing phase will only be determined by the new location of the animal in the place field. Therefore, the firing phase in the first bump oscillation cycle after the network perturbation is more advanced than the firing phase in the last bump oscillation cycle right before the perturbation, and the amount of precession is similar to that in the case without perturbation (Fig. 6e). This agrees well with the experimental observation (Fig. 6b), and indicates that the phase-position code is robust to the perturbation of the hippocampal dynamics.

Overall, our model reproduces these two experimental findings, and suggests that there exists a one-to-one correspondence between the firing phase of a place cell and the travelled distance in the neuron’s place field, which is independent of the animal’s running speed or the perturbation duration (Fig. S4). This agrees well with experimental observations (***O’Keefe and Recce, 1993***) that theta phase correlates better with the animal’s location than with time (Fig. 6f&g). In addition to the results for unimodal cells as introduced above, our model predicts new results for bimodal cells. First, in contrast to a unimodal cell, a bimodal cell will have two peaks in its firing frequency, with one slightly higher than the LFP theta baseline (due to phase precession) and the other slightly lower than the LFP theta baseline (due to phase procession). The precession-associated frequency positively correlates with the running speed of the animal, while the procession-associated frequency negatively correlates with the running speed (Fig. 6d). Second, similar to the preserved phase shift in unimodal cells, both the phase precession and procession of a bimodal cell after transient intrahippocampal perturbation continue from the new location of the animal (see Fig. S5), no matter how long the silencing period lasts. The two predictions could be tested by experiments.

## Discussion

### Model contributions

In this paper, we have proposed a CANN with firing rate adaptation to unveil the underlying mechanism of place cell phase shift during locomotion. We show that the interplay between intrinsic mobility (owing to firing rate adaptation) and extrinsic mobility (owing to the location-dependent sensory inputs) leads to an oscillatory tracking state, which naturally accounts for theta sweeps where the decoded position oscillates around the animal’s physical location at the theta rhythm. At the single neuron level, we show that the forward and backward bump sweeps account for, respectively, phase precession and phase procession. Furthermore, we show that the varied adaptation strength explains the emergence of bimodal and unimodal cells, that is, as the adaptation strength increases, forward sweeps of the bump gradually suppress backward sweeps, and as a result, neurons initially exhibiting both significant phase precession and procession (due to a low level adaptation) will gradually exhibit only predominant phase precession (due to a high level adaptation).

### Computational models for theta phase shift and theta sweeps

As a subject of network dynamics, oscillatory tracking has been studied previously in an excitatory-inhibitory neural network (***Folias and Bressloff, 2004***), where it was found that decreasing the external input strength can lead to periodic emission of traveling waves in the network (Hopf instability), which is analogous to the oscillatory tracking state in our model. However, their focus was on the mathematical analysis of such dynamical behavior, while our focus is on the biological implications of oscillatory tracking, i.e., how can it be linked to phase precession and procession of hippocampal place cells.

Due to their potential contributions to the temporal sequence learning involved in spatial navigation and episodic memory (***Mehta et al., 1997, 2002***; ***Yamaguchi, 2003***), theta phase precession and forward theta sweeps have been modelled in the field for decades. These models can be divided into two main categories, with one relying on the mechanism of single cell oscillation (***O’Keefe and Recce, 1993***; ***Kamondi et al., 1998***; ***Lengyel et al., 2003***; ***O’keefe and Burgess, 2005***; ***Mehta et al., 2002***), and the other relying on the mechanism of recurrent interactions between neurons (***Tsodyks et al., 1996***; ***Romani and Tsodyks, 2015***; ***Kang and DeWeese, 2019***). A representative example of the former is the oscillatory interference model (***O’Keefe and Recce, 1993***; ***Lengyel et al., 2003***), which produces phase precession via the superposition of two oscillatory signals, with one from the baseline somatic oscillation at the LFP theta frequency (reflecting the inputs from the medial septal pacemaker (***Stewart and Fox, 1990***)), and the other from the dendritic oscillation whose frequency is slightly higher. While these models can explain a large variety of experimental phenomena, it remain unclear how oscillation of individual neurons has a frequency higher than the baseline theta frequency. Here, our model provides a network mechanism for how such higher-frequency oscillation emerges.

A representative model relying on neuronal recurrent interactions is the activation spreading model (***Tsodyks et al., 1996***). This model produces phase precession via the propagation of neural activity along the movement direction, which relies on asymmetric synaptic connections. A later version of this model considers short-term synaptic plasticity (short-term depression) to implicitly implement asymmetric connections between place cells (***Romani and Tsodyks, 2015***), and reproduces many other interesting phenomena, such as phase precession in different environments. Different from these two models, our model considers firing rate adaptation to implement symmetry breaking and hence generates activity propagation. To prevent the activity bump from spreading away, their model considers an external theta input to reset the bump location at the end of each theta cycle, whereas our model generates an internal oscillatory state, where the activity bump travels back due to the attraction of external location input once it spreads too far away. Moreover, theoretical analysis of our model reveals how the adaptation strength affect the direction of theta sweeps, as well as offers a more detailed understanding of theta cycling in complex environments.

Based on our simulation, both STD and SFA show the ability to produce bi-directional sweeps within a CANN model, with the SFA uniquely enabling uni-directional sweeps in the absence of external theta inputs. This difference might be due to the lack of exhaustively exploration of the entire parameter space. However, it might also attribute to the subtle yet important theoretical distinctions between STD and SFA. Specifically, STD attenuates the neural activity through a reduction in recurrent connection strength, whereas SFA provides inhibitory input directly to the neurons, potentially impacting all excitatory inputs. These differences might explain the diverse dynamical behaviors observed in our simulations. Future experiments could clarify these distinctions by monitoring changes in synaptic strength and inhibitory channel activation during theta sweeps.

### Beyond the linear track environment

Besides the linear track environment, the mechanism of generating theta sweeps proposed in our model can also be generalized to more complex environments. For instance, in a T-maze environment, our model explains the constant cycling of theta sweeps between left and right arms. Such cycling behavior may be important for high-speed actions such as predating and escaping which require animals to make decision among several future scenarios at the sub-second level. Similar alternative activity sweeps in the T-maze environment has been studied in a previous paper (***Romani and Tsodyks, 2015***), which showed that the frequency of alternation correlates with overtly deliberative behaviors such as head scans (frequency at 1 Hz or less) (***Johnson and Redish, 2007***). In contrast to our model, the network activity in their model propagates continuously from the current location on the center arm till the end of the outer arm, which takes a few theta cycles (i.e., 1 second or more). In our model, the network bump alternately sweeps to one of the two outer arms at a much higher frequency (∼ 8 Hz), which may be related to fast decision-making or planing in natural environments (***Kay et al., 2020***). Furthermore, our model can also be easily extended to the multiple-arms (> 2) environment (***Gillespie et al., 2021***) or the cascade-T environment (***Johnson and Redish, 2007***) with the underlying mechanism of generating theta cycling remaining unchanged. In addition to the linear and T-maze environments, phase shift has also been reported when an animal navigates in an open field environment. However, due to the lack of recorded neurons, decoding theta sweeps in the 2D environment is not as straightforward as in the 1D case. While theta sweeps in the 1D case have been associated with goal-directed behaviors and spatial planning (***Wikenheiser and Redish, 2015***), it remains unclear whether such conclusion is applicable to the 2D case. Our preliminary result shows that in the 2D CANN where neurons are arranged homogeneously according to their relative firing locations, the activity bump will sweep along the tangent direction of the movement trajectory, similar to the 1D case (see SI.4 and Fig. S6 for details). It will be interesting to explore theta sweeps in the open field environment in detail when more experimental data is available.

### Model predictions and future works

Our model has several predictions which can be tested in future experiments. For instance, the height of the activity bump in the forward sweep window is higher than that in the backward sweep window (Fig. 4c) due to the asymmetric suppression effect from the adaptation. For bimodal cells, they will have two peaks in their firing frequency as the animal runs across the firing fields, with one corresponding to phase precession and the other corresponding to phase procession. Similar to unimodal cells, both the phase precession and procession of a bimodal cell after transient intrahippocampal perturbation will continue from the new location of the animal (Fig. S5). Interestingly, our model of the T-maze environment showed an expected phenomenon that as the animal runs towards the decision point, the theta sweep length also shows cyclical patterns (Fig. 5b lower panel). An intuitive explanation is that, due to the slow dynamics in firing rate adaptation (with a large time constant compared to neural firing), a long sweep leads to an adaptation effect on the neurons at the end of the sweep path. Consequently, the activity bump cannot travel as far due to the adaptation effect on those neurons, resulting in a shorter sweep length compared to the previous one. In the next round, the activity bump exhibits a longer sweep again because those neurons have recovered from the previous adaptation effect. We plan to test this phenomenon in future experiments.

In the current study, we have modeled the place-cell population in the hippocampus with a CANN and adopted firing rate adaptation to generate theta phase shift. In fact, this model can be easily extended to the grid cell population without changing the underlying mechanism. For instance, we can induce the torus-like connection profile (periodic boundary in the 2D space) (***Samsonovich and McNaughton, 1997***; ***McNaughton et al., 2006***) or the locally inhibitory connection profile (***Burak and Fiete, 2009***; ***Couey et al., 2013***) in the CANN structure to construct a grid cell model, and by imposing firing rate adaptation, neurons in the grid cell network will also exhibit phase shift as the animal moves through the grid field, as reported in previous experimental studies (***Hafting et al., 2008***; ***Van Der Meer and Redish, 2011***). Notably, although for both grid cells and place cells, CANNs can generate theta phase shift, it does not mean that they are independent from each other. Instead, they might be coordinated by the same external input from the environment, as well as by the medial septum which is known to be a pacemaker that synchronises theta oscillations across different brain regions (***King et al., 1998***; ***Wang, 2002***). We will investigate this issue in future work.

Our model also suggests that the “online” theta sweep and the “offline” replay may share some common features in their underlying mechanisms (***Romani and Tsodyks, 2015***; ***Hopfield, 2010***; ***Kang and DeWeese, 2019***; ***Jahnke et al., 2015***). We have shown that the activity bump with strong adaptation can move spontaneously when the external input becomes weak enough (see previous sections). Such non-local spreading of neural activity has a speed much faster than the conventional speed of animals (the external input speed in our model, see Fig. 2d), which resembles the fast spreading of the decoded position during sharp-wave ripple events (***Diba and Buzsáki, 2007***; ***Foster and Wilson, 2006***; ***Karlsson and Frank, 2009***; ***Dragoi and Tonegawa, 2011***). This indicates that these two phenomena may be generated by the same neural mechanism of firing rate adaptation, with theta sweeps originating from the interplay between the adaptation and the external input, while replay originating from only the adaptation, since the external input is relatively weak during the “offline” state. This hypothesis seems to be supported by the coordinated emergence of theta sequences and replays during the post-natal development period (***Muessig et al., 2019***), as well as their simultaneous degradation when the animal travelled passively on a model train (***Drieu et al., 2018***).

Nevertheless, it is important to note that the CANN we adopt in the current study is an idealized model for the place cell population, where many biological details are missed (***Amari, 1977***; ***Tsodyks and Sejnowski, 1995***; ***Samsonovich and McNaughton, 1997***; ***Tsodyks, 1999***). For instance, we have assumed that neuronal synaptic connections are translation-invariant in the space. In practice, such a connection pattern may be learned by a synaptic plasticity rule at the behavioral time scale when the animal navigates actively in the environment (***Bittner et al., 2017***). In future work, we will explore the detailed implementation of this connection pattern, as well as other biological correspondences of our idealized model, to establish a comprehensive picture of how theta phase shift is generated in the brain.

## Materials and Methods

### General summary of the model

We consider a one-dimensional continuous attractor neural network (1D CANN), in which neurons are uniformly aligned according to their firing fields on a linear track (for the T-maze case, see Methods. below; for the case of the open field (2D CANN), see SI.4). Denote *U (x, t*) the synaptic input received by the place cell at location *x*, and *r*(*x, t)* the corresponding firing rate. The dynamics of the network is written as

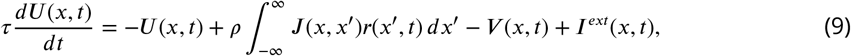

where *τ* is the time constant of *U (x, t*) and *ρ* the neuron density. The firing rate *r*(*x, t)* is given by

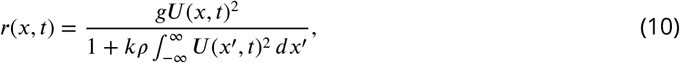

where *k* controls the strength of the global inhibition (divisive normalization), *g*denotes a gain factor. *J* (*x, x*^′^) denotes the connection weight between place cells at location *x* and *x*^′^, which is written as:

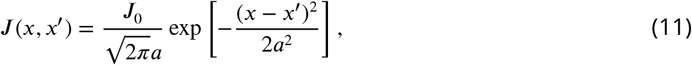

where *J*_0_ controls the strength of the recurrent connection and *a* the range of neuronal interaction. Notably, *J* (*x, x*^′^) depends on the relative distance between two neurons, rather than the absolute locations of neurons. Such translation-invariant connection form is crucial for the neutral stability of the attractor states of CANNs (***Wu et al., 2016***). *I*^*ext*^*(x, t*) represents the external input which conveys the animal location information to the hippocampal network, which is written as:

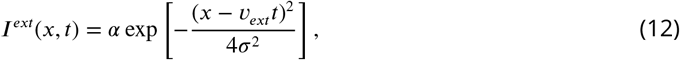

with *v*_*ext*_ denoting the animal’s running speed and *α* controlling the input strength to the hippocampus. *σ* denotes the width of the external input *I*^*ext*^, which is set to be equal to the recurrent connection width *a* in the main text and the following derivation. *V (x, t*) denotes the adaptation effect of the place cell at location *x*, which increases with the synaptic input (and hence the place cell’s firing rate), i.e.,

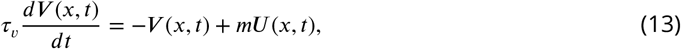

with *τ*_*v*_ denoting the time constant of *V (x, t*) and *m* the adaptation strength. Note that *τ*_*v*_ ≫ *τ*, meaning that adaptation is a much slower process compared to the neural firing.

### Stability analysis of the bump state

We derive the condition under which the bump activity is the stable state of the CANN. For simplicity, we consider the simplest case that there is no external input and adaptation in the network, i.e., *m* = *α* = 0. In this case, the network state is determined by the strength of the recurrent excitation and global inhibition. When the global inhibition is strong (*k* is large), the network is silent, i.e., no bump activity emerges in the CANN. When the global inhibition is small, an activity bump with the Gaussian-shaped profile emerges, which is written as:

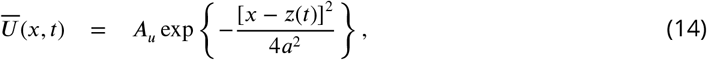

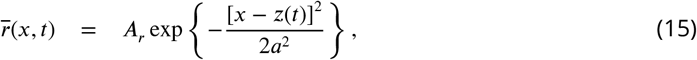

with *A*_*u*_ and *A*_*r*_ representing the amplitudes of the synaptic input bump and the firing rate bump, respectively. *z*(*t*) represents the bump center, and *a* is the range of neuronal interaction (defined in Methods. ). To solve the network dynamics, we substitute Eqs. 14&15 into Eqs. 9&10, which gives (see SI.2 for more details of the derivation):

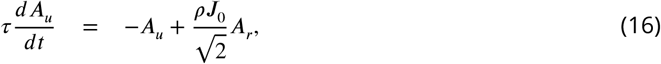

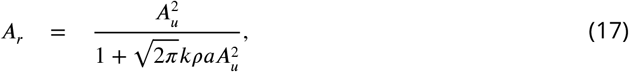

These two equations describes how the bump amplitudes change with time. For instance, if neurons are weakly connected (small *J*_0_) or they are connected sparsely (small *ρ*), the second term on the right-hand side of Eq. 16 is small, and *A*_*u*_ will decay to zero, implying that the CANN cannot sustain a bump activity. By setting *dA*_*u*_*/dt* = 0, we obtain:

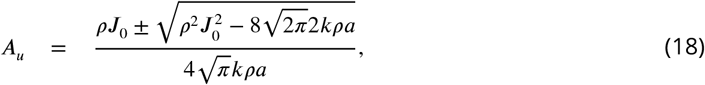

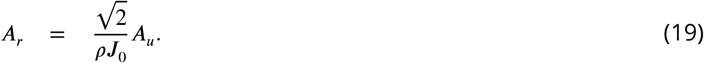

It is straightforward to check that only when:

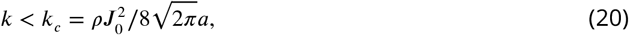

*A*_*u*_ have two real solutions (indicated by the ± sign in Eq. 18), i.e., the dynamic system (Eqs. 16&17) has two fixed points. It can be checked that only 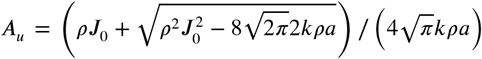 is the stable solution.

### Analysis of the intrinsic mobility of the bump state

We derive the condition under which the bump of the CANN moves spontaneously in the attractor space without relying on external inputs. As the adaptation strength increases, the bump activity becomes unstable and has tendency to move away from its location spontaneously. Such intrinsic mobility of the CANN has been shown in previous studies (***Bressloff, 2011***; ***Wu et al., 2016***; ***Mi et al., 2014***). We set *α* = 0 (no external input), and investigate the effect of adaptation strength *m* on the bump dynamics. Our simulation result shows that during the spontaneous movement, *V (x, t*) can also be represented by a Gaussian-shaped bump, which is written as

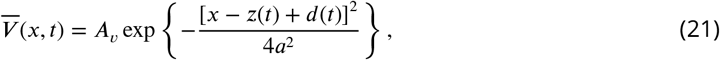

where *A*_*v*_ denotes the amplitude of the adaptation bump, and *d*(*t*) the displacement between the bump centers of *U (x, t*) and *V (x, t*). This displacement originates from the slow dynamics of adaptation, which leads to the adaptation bump always lags behind the neural activity bump. Similar to Methods., we substitute the bump profiles Eqs. (14, 15, 21) into the network dynamics Eqs. (9, 10, 13), and obtain:

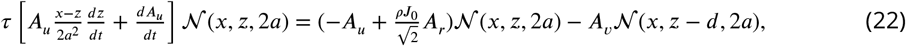

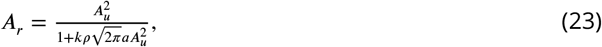

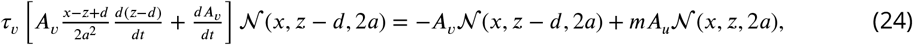

where 𝒩 (*x, z*, 2*a*) = exp{− [*x − z*]^2^/4*a*^2^}.

At first glance, the resulting equations given by Eqs. (22) and (24) may seem intractable due to the high dimensionality (i.e., 2*N*, where *N* is the number of neurons in the network). However, a key property of CANNs is that their dynamics are dominated by a few motion modes, which correspond to distortions of the bump shape in terms of height, position, width, etc. ***Fung et al. (2010***). By projecting the network dynamics onto its dominant motion modes ***Fung et al. (2010***) (which involves computing the inner product of a function *f(x)* with a mode *u*_*n*_*(x))*, we can significantly simplify the network dynamics. Typically, projecting onto the first two motion modes is sufficient to capture the main features of the dynamics, which are given by,

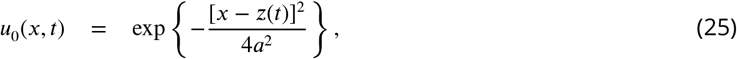

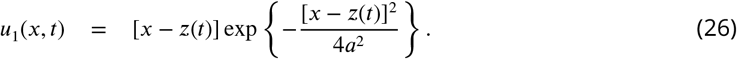

By projecting the network dynamics onto these two motion modes, we obtain:

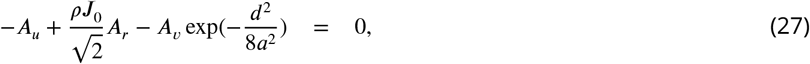

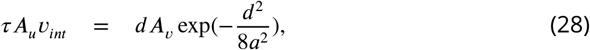

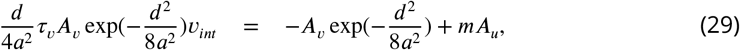

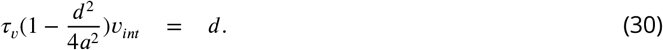

Note that we assume that the bump height keep as constant over time, i.e., *dA*_*u*_*/dt* = *dA*_*v*_*/dt* = 0 is assumed. Eqs. 27-30 describes the relationships between bump features *A*_*u*_, *A*_*r*_, *A*_*v*_, *v*_*int*_ and *d*, where *v*_*int*_ *= dz/dt* representing the intrinsic moving speed of the bump center. By solving these equations together with eqn. 23, we obtain,

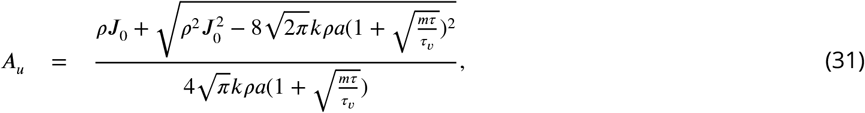

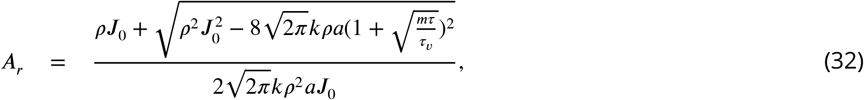

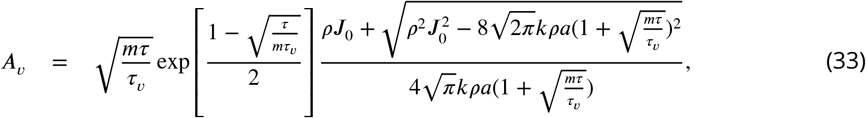

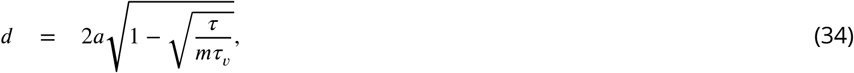

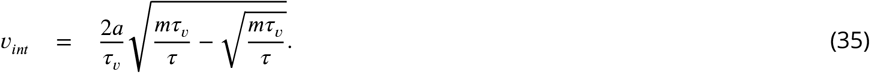

Eqs. 31-33 describe the amplitudes of the bumps of synaptic input, firing rate, and adaptation in the CANN, respectively, and Eq. 34 describes the displacement between the neural activity and adaptation bumps. From Eq. 35, we see that for the bump to travel spontaneously, it requires *m* > *τ/τ*_*v*_, i.e., the adaptation strength is larger than a threshold given by the ratio between two time constants *τ* and *τ*_*v*_. As the adaptation strength increases (larger *m*), the travelling speed of the bump increases (larger *v*_*int*_).

### Analysis of the oscillatory tracking behaviour of the bump state

When both the external input and the adaptation are applied to the CANN, the bump activity can oscillate around the external input if the strengths of the external input and the adaptation are appropriated. The simulation shows that during the oscillatory tracking, the bump shape is roughly unchanged, and the oscillation of the bump center can be approximated as a sinusoidal wave expressed as:

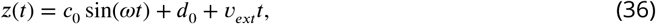

where *c*_0_ and *ω*denote, respectively, the oscillation amplitude and frequency, and *d*_0_ denotes a constant offset between the oscillation center and the external input.

Similar to the analysis in Methods., we substitute the expression of *z*(*t*) (Eq. 36) into Eqs. (14, 15, 21), and then simplify the network dynamics by applying the projection method (see SI.3 for more detailed derivation). We obtain,

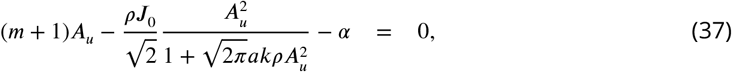

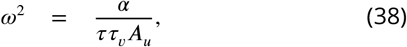

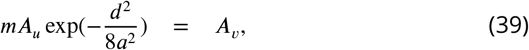

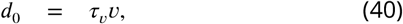

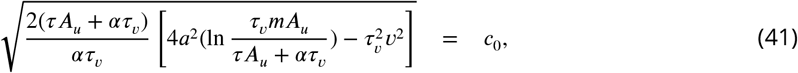

Eqs. 37-41 describe the relationships among 6 oscillation features *A*_*u*_, *A*_*r*_, *A*_*v*_, *c*_0_, *d*_0_ and *ω*. By solving these equations, we obtain:

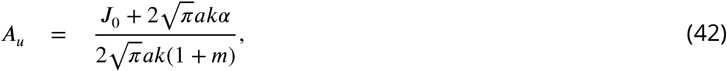

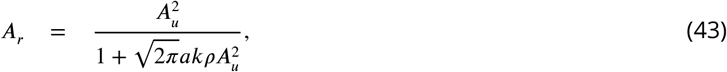

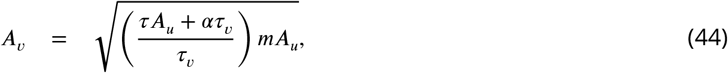

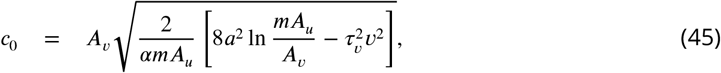

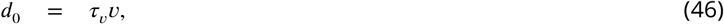

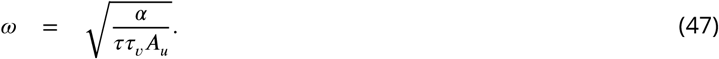

It can be seen from Eq. 45 that for the bump activity to oscillate around the external input (i.e., the oscillation amplitude *c*_0_ > 0), it requires that 8*a*^2^ ln 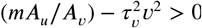 . This condition gives the boundary (on the parameter values of the input strength *α* and the adaptation strength *m*) that separate two tracking states, i.e., smooth tracking and oscillatory tracking (see Fig. 2g and Fig. S2 for the comparison between the simulation results and theoretical results).

Note that to get the results in Eqs. 37-41, we have assumed that the amplitudes of neural activity bumps and the adaptation bump remain unchanged during the oscillation (i.e., *A*_*u*_, *A*_*v*_, *A*_*r*_ are constants). However, this assumption is not satisfied when the SFA strength *m* is large (see previous sections and Fig. 4). In such a case, we carry out simulation to analyze the network dynamics.

### Implementation details of the linear track environment

For the linear track environment, we simulate an 1D CANN with 512 place cells topographically organized on the one-dimensional neuronal track. Since we are interested in how the neuronal firing phase shifts as the animal moves through the firing field of a place cell, we investigate the place cell at location *x* = 0 and ignore the boundary effect, that is, we treat the linear track with the infinite length. The neural firing time constant is set to be 3 ms, while the time constant of spike frequency adaptation is much longer, which is set to be 144 ms. The density of place cells on the linear track is set to be 256*/π.*The excitatory interaction range of place cells is set to be 0.4*m*, while the maximum excitatory connection strength *J*_0_ is set to be 0.2. The gain factor is set to be 5. The global inhibition strength *k* is set to be 5. The moving speed of the virtual animal *v*_*ext*_ is set to be 1.5 m/s. For the simulation details, we use the first-order Euler method with the time step *δt* set to be 0.3 the duration of simulation *T* set to be 10 s. These parameters are commonly used in all plots related to the linear track environment (see Table.1 for a summary).

**Table 1.** Commonly used parameter values in the simulation of the linear track environment.

| Parameters | Values |
| --- | --- |
| Number of place cells: $N$ | 512 |
| Time constant of neural firing: $\tau$ | 3ms |
| Time constant of spike frequency adaptation: $\tau_v$ | 144ms |
| Neuron density: $\rho$ | $256/\pi$ |
| Recurrent connection range (Gaussian width): $a$ | 0.4 m |
| Width of external input (Gaussian width): $\sigma$ | 0.4 m |
| Recurrent connection strength: $J_0$ | 0.2 |
| Gain factor: $g$ | 5 |
| Global inhibition strength: $k$ | 5 |
| Moving speed of the external input: $v_{ext}$ (m/s) | 1.5 |
| Time interval: $\delta t$ | 0.3 s |
| Simulation duration: $T$ | 10 s |

For the two key parameters, i.e., the external input strength *α* and the adaptation strength *m*, we vary their values in different plots. Specifically, for illustrating the smooth tracking state in Fig. 2c, we set *α* = 0.19 and *m* = 0. For illustrating the travelling wave state (intrinsic mobility of the bump state) in Fig. 2d, we set *α* = 0 and *m* = 0.31. For plotting the relationship between the intrinsic speed *v*_*int*_ and the adaptation strength *m* shown in Fig. 2e, we keep *α* = 0, but vary *m* in the range between 0 and 0.1 with a step of 0.05. For plotting the overall phase diagram including all three moving states as shown in Fig. 2g, we vary *α* in the range between 0.05 and 0.16 with a step of 0.001, and *m* in the range between 0.9 and 1.8 with a step of 0.01. To generate bimodal cell firing patterns in Fig. 3a and Fig. 4a,e&g, we choose *α* = 0.19 and *m* = 3.02. To generate unimodal firing patterns in Fig. 4b,f&h, we choose *α* = 0.19 but a relatively larger adaptation strength with *m* = 3.125. The values of these two parameters in different plots are summarized in Table. 2.

**Table 2.** Figure specific parameter values for input strength *α* and adaptation strength *m*.

| Figures/parameters | $\alpha$ | $m$ |
| --- | --- | --- |
| An example of smooth tracking (Fig. 2c) | 0.19 | 0 |
| An example of traveling wave (Fig. 2d) | 0 | 0.31 |
| Intrinsic speed vs. adaptation strength (Fig. 2e) | 0 | 0:0.05:0.1 |
| Phase diagram (Fig. 2g) | 0.05:0.001:0.16 | 0.9:0.01:1.8 |
| Oscillatory tracking (bimodal) (Fig. 4a,e,g) | 0.19 | 3.02 |
| Oscillatory tracking (unimodal) (Fig. 4b,f,h) | 0.19 | 3.125 |

### Implementation details of the T-maze environment

#### Parameter configurations during simulation

To simulate the T-maze environment, we consider a CANN in which place cells are topographically organized in a T-shaped area which consists of a vertical central arm and two horizontal left and right arms (Fig. 5a). The width of the central arm is set to be 0.84 m and the length is set to be 3.14 m. The widths of the two horizontal arms are also set to be 0.84 m, while the lengths of both arms are set to be 2.36 m. The connection strength between two neurons is determined by the distance between them, which is written as:

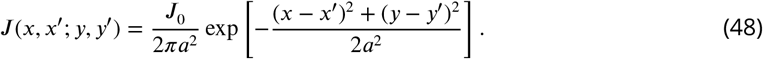

Here (*x, y)* and (*x*^′^, *y*^′^) represent the coordinates of two neurons in the T-maze environment, *a* is the recurrent connection range which is set to be 0.3, and *J*_0_ controls the connection strength which is set to be 0.0125. Since we are interested in investigating theta sweeps when the animal is running on the central arm towards the junction point, the external input is restricted on the central arm which is modelled by a Gaussian-like moving bump written as:

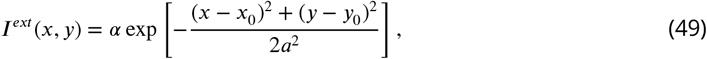

where *x*_0_ = 0 and *y*_0_ *= v*_*ext*_*t*represent the center location of the external input with a moving speed *v*_*ext*_ = 1.5 m/s. In the simulation, we used the first-order Euler method with the time step *δt* = 0.3 s and the duration of simulation *T* = 4.2*s*. The parameters used are summarized in Table.3.

**Table 3.**
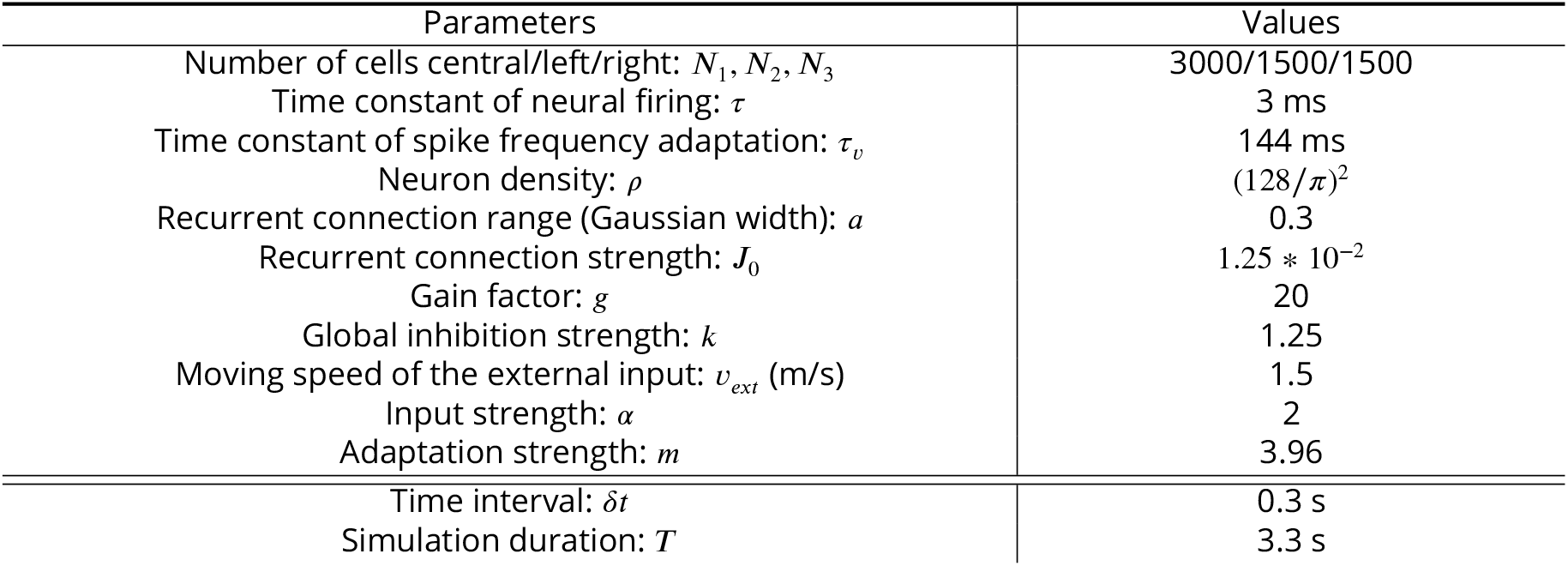
Parameters values in the simulation of the T-maze environment.

#### Calculating auto-correlogram and cross-correlogram

To show the “cycle skipping” effect of a single place cell in the T-maze environment, we calculate the auto-correlogram of the firing rate trace of a place cell whose firing field encodes a location on the left arm (the upper panel in Fig. 5d). Assume the firing trace of the place cell is *f (t)* (showed in left panel in Fig. 5c), the auto-correlogram is calculated as:

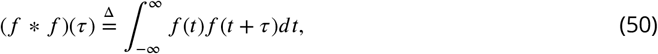

where *τ* represents the time offset.

To show the “alternative cycling” effect of a pair of place cells with each of them encoding a location on each of the two outward arms, we calculate the cross-correlogram between their firing traces (the lower panel in Fig. 5d). It measures the similarity of the two firing traces as a function of the temporal offset of one relative to the other. Assume the firing traces of the two place cells are *f (t)* and *g(t)*, respectively, the cross-correlogram is calculated as:

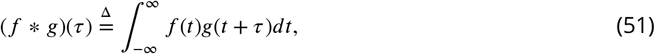

where *τ* represents the time offset.

### Details of generating the probability heatmap of theta phase shift

In Fig. 4g&h we described the smoothed probability heatmaps of theta phase versus normalized position in the place field of both bimodal and unimodal cells. Generally, these two plots are similar to the traditional spike plot of phase and position traveled in the place field (***O’Keefe and Recce, 1993***; ***Skaggs et al., 1996***). However, in our rate-based model, the phase of neuronal spike is not directly modelled, rather we use the phase of firing rate peak to represent the phase shift in neuronal firing. Here we describe the implementation details of generating the heatmaps.

The x-axis denotes the normalized position in the place field, with -1 representing the position where the animal just enters the place field, and 1 representing the position where the animal just leaves the place field. In our simulation, the firing field of a place cell with preferred location at *x*_0_ is defined as *x ∈ (x*_0_ − 2.5 *\* a, x*_0_ + 2.5*a*), with *a* roughly the half size of the firing field. Consider the animal is at *x*_*t*_ *at* time *t* (note that *x*_*t*_ *= v*_*ext*_*t)*, then its normalized position 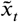 is calculated as 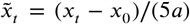 . The y-axis represents the phase of neuronal activity, which is in the range of (0°, 720°). To calculate the phase at every time step, we divide the duration of the animal traversing the linear track into multiple theta cycles according to the bump’s oscillation. We can calculate the phase by *θ*_*t*_ *= (t − t*_0_*)/T*, with *t*_0_ referring to the beginning of the present theta cycle and T referring to the theta period. Denote the firing rate of the *i*-th neuron at time t as 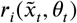, the probability heatmap is calculated by,

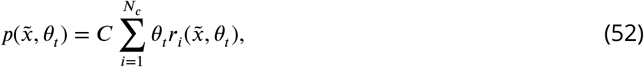

where 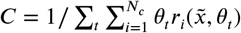 is the normalization factor.

### Spike generation from the firing rate

To understand phase shift based on spiking time rather than the peak firing rate, we convert the firing rate into spike trains according to the Poisson statistics (note that our analysis is rate-based, but converting to spike-based does not change the underlying mechanism). For the ith place cell which encodes position *x*_*i*_ on the linear track, the number of spikes *n*_*i*_ it generates within a time interval Δ*t* satisfies a Poisson distribution, which is expressed as,

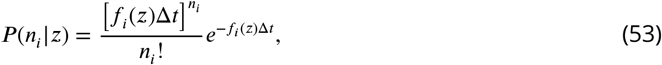

where *z* is the animal’s location, and *f*_*i*_*(z)* is the tuning function of cell *i,*which is given by

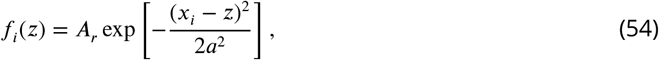

where *A*_*r*_ denotes the amplitude of the neural activity bump and *a* the range of recurrent interaction.

## Acknowledgements

We thank Brad Pfeiffer for sharing the data. We also thank Brad Pheiffer, Cheng Wang, Li Yao for valuable discussions.

## Author Contributions

TC, ZJ and SW conceptualized and designed the research. TC, ZJ, JZ, and YM analyzed the model and performed the simulations. ZJ, WZ, TH, NB and SW interpreted the results. ZJ, TC, DB, NB and SW wrote the manuscript.

## Data and materials availability

All code for reproducing the figures in the main text are available in the supplementary materials.

## List of material contained in the Supplementary Material

- Supplementary text (pdf file)
- Figures S1-S4
- Video 1-4
- Code for reproducing all the figures in the main text

## References

Alonso, A. and Klink, R. (1993). Differential electroresponsiveness of stellate and pyramidal-like cells of medial entorhinal cortex layer ii. Journal of neurophysiology, 70(1):128–143.

Amari, S.-i. (1977). Dynamics of pattern formation in lateral-inhibition type neural fields. Biological cybernetics, 27(2):77–87.

Azizi, A. H., Wiskott, L., and Cheng, S. (2013). A computational model for preplay in the hippocampus. Frontiers in computational neuroscience, 7:161.

Battaglia, F. P. and Treves, A. (1998). Attractor neural networks storing multiple space representations: a model for hippocampal place fields. Physical Review E, 58(6):7738.

Benda, J. and Herz, A. V. (2003). A universal model for spike-frequency adaptation. Neural computation, 15(11):2523–2564.

Bittner, K. C., Milstein, A. D., Grienberger, C., Romani, S., and Magee, J. C. (2017). Behavioral time scale synaptic plasticity underlies ca1 place fields. Science, 357(6355):1033–1036.

Brandon, M. P., Bogaard, A. R., Schultheiss, N. W., and Hasselmo, M. E. (2013). Segregation of cortical head direction cell assemblies on alternating theta cycles. Nature neuroscience, 16(6):739–748.

Bressloff, P. C. (2011). Spatiotemporal dynamics of continuum neural fields. Journal of Physics A: Mathematical and Theoretical, 45(3):033001.

Burak, Y. and Fiete, I. R. (2009). Accurate path integration in continuous attractor network models of grid cells. PLoS computational biology, 5(2):e1000291.

Burgess, N., Recce, M., and O’Keefe, J. (1994). A model of hippocampal function. Neural networks, 7(6-7):1065–1081.

Couey, J. J., Witoelar, A., Zhang, S.-J., Zheng, K., Ye, J., Dunn, B., Czajkowski, R., Moser, M.-B., Moser, E. I., Roudi, Y., et al. (2013). Recurrent inhibitory circuitry as a mechanism for grid formation. Nature neuroscience, 16(3):318–324.

Deshmukh, S. S., Yoganarasimha, D., Voicu, H., and Knierim, J. J. (2010). Theta modulation in the medial and the lateral entorhinal cortices. Journal of neurophysiology, 104(2):994–1006.

Diba, K. and Buzsáki, G. (2007). Forward and reverse hippocampal place-cell sequences during ripples. Nature neuroscience, 10(10):1241–1242.

Dong, X., Chu, T., Huang, T., Ji, Z., and Wu, S. (2021). Noisy adaptation generates lévy flights in attractor neural networks. Advances in Neural Information Processing Systems, 34:16791–16804.

Dragoi, G. and Tonegawa, S. (2011). Preplay of future place cell sequences by hippocampal cellular assemblies. Nature, 469(7330):397–401.

Drieu, C., Todorova, R., and Zugaro, M. (2018). Nested sequences of hippocampal assemblies during behavior support subsequent sleep replay. Science, 362(6415):675–679.

Feng, T., Silva, D., and Foster, D. J. (2015). Dissociation between the experience-dependent development of hippocampal theta sequences and single-trial phase precession. Journal of Neuroscience, 35(12):4890–4902.

Fenton, A. A. and Muller, R. U. (1998). Place cell discharge is extremely variable during individual passes of the rat through the firing field. Proceedings of the National Academy of Sciences, 95(6):3182–3187.

Fernández-Ruiz, A., Oliva, A., Nagy, G. A., Maurer, A. P., Berényi, A., and Buzsáki, G. (2017). Entorhinal-ca3 dual-input control of spike timing in the hippocampus by theta-gamma coupling. Neuron, 93(5):1213–1226.

Folias, S. E. and Bressloff, P. C. (2004). Breathing pulses in an excitatory neural network. SIAM Journal on Applied Dynamical Systems, 3(3):378–407.

Foster, D. J., Morris, R. G., and Dayan, P. (2000). A model of hippocampally dependent navigation, using the temporal difference learning rule. Hippocampus, 10(1):1–16.

Foster, D. J. and Wilson, M. A. (2006). Reverse replay of behavioural sequences in hippocampal place cells during the awake state. Nature, 440(7084):680–683.

Foster, D. J. and Wilson, M. A. (2007). Hippocampal theta sequences. Hippocampus, 17(11):1093–1099.

Fuhrmann, G., Markram, H., and Tsodyks, M. (2002). Spike frequency adaptation and neocortical rhythms. Journal of neurophysiology, 88(2):761–770.

Fung, C. A., Wong, K. M., and Wu, S. (2010). A moving bump in a continuous manifold: a comprehensive study of the tracking dynamics of continuous attractor neural networks. Neural Computation, 22(3):752–792.

Geisler, C., Robbe, D., Zugaro, M., Sirota, A., and Buzsáki, G. (2007). Hippocampal place cell assemblies are speed-controlled oscillators. Proceedings of the National Academy of Sciences, 104(19):8149–8154.

Gillespie, A. K., Maya, D. A. A., Denovellis, E. L., Liu, D. F., Kastner, D. B., Coulter, M. E., Roumis, D. K., Eden, U. T., and Frank, L. M. (2021). Hippocampal replay reflects specific past experiences rather than a plan for subsequent choice. Neuron, 109(19):3149–3163.

Hafting, T., Fyhn, M., Bonnevie, T., Moser, M.-B., and Moser, E. I. (2008). Hippocampus-independent phase precession in entorhinal grid cells. Nature, 453(7199):1248–1252.

Hao, J., Wang, X.-d., Dan, Y., Poo, M.-m., and Zhang, X.-h. (2009). An arithmetic rule for spatial summation of excitatory and inhibitory inputs in pyramidal neurons. Proceedings of the National Academy of Sciences, 106(51):21906–21911.

Harris, K. D., Henze, D. A., Hirase, H., Leinekugel, X., Dragoi, G., Czurkó, A., and Buzsáki, G. (2002). Spike train dynamics predicts theta-related phase precession in hippocampal pyramidal cells. Nature, 417(6890):738– 741.

Hassabis, D., Kumaran, D., Vann, S. D., and Maguire, E. A. (2007). Patients with hippocampal amnesia cannot imagine new experiences. Proceedings of the National Academy of Sciences, 104(5):1726–1731.

Hopfield, J. J. (2010). Neurodynamics of mental exploration. Proceedings of the National Academy of Sciences, 107(4):1648–1653.

Jahnke, S., Timme, M., and Memmesheimer, R.-M. (2015). A unified dynamic model for learning, replay, and sharp-wave/ripples. Journal of Neuroscience, 35(49):16236–16258.

Jaramillo, J. and Kempter, R. (2017). Phase precession: a neural code underlying episodic memory? Current opinion in neurobiology, 43:130–138.

Jensen, O. and Lisman, J. E. (2000). Position reconstruction from an ensemble of hippocampal place cells: contribution of theta phase coding. Journal of neurophysiology, 83(5):2602–2609.

Johnson, A. and Redish, A. D. (2007). Neural ensembles in ca3 transiently encode paths forward of the animal at a decision point. Journal of Neuroscience, 27(45):12176–12189.

Kamondi, A., Acsády, L., Wang, X.-J., and Buzsáki, G. (1998). Theta oscillations in somata and dendrites of hippocampal pyramidal cells in vivo: Activity-dependent phase-precession of action potentials. Hippocampus, 8(3):244–261.

Kang, L. and DeWeese, M. R. (2019). Replay as wavefronts and theta sequences as bump oscillations in a grid cell attractor network. Elife, 8:e46351.

Karlsson, M. P. and Frank, L. M. (2009). Awake replay of remote experiences in the hippocampus. Nature neuroscience, 12(7):913–918.

Kay, K., Chung, J. E., Sosa, M., Schor, J. S., Karlsson, M. P., Larkin, M. C., Liu, D. F., and Frank, L. M. (2020). Constant sub-second cycling between representations of possible futures in the hippocampus. Cell, 180(3):552–567.

King, C., Recce, M., and O’keefe, J. (1998). The rhythmicity of cells of the medial septum/diagonal band of broca in the awake freely moving rat: relationships with behaviour and hippocampal theta. European Journal of Neuroscience, 10(2):464–477.

Lengyel, M., Szatmáry, Z., and Érdi, P. (2003). Dynamically detuned oscillations account for the coupled rate and temporal code of place cell firing. Hippocampus, 13(6):700–714.

Losonczy, A., Zemelman, B. V., Vaziri, A., and Magee, J. C. (2010). Network mechanisms of theta related neuronal activity in hippocampal ca1 pyramidal neurons. Nature neuroscience, 13(8):967–972.

McNaughton, B. L., Battaglia, F. P., Jensen, O., Moser, E. I., and Moser, M.-B. (2006). Path integration and the neural basis of the’cognitive map’. Nature Reviews Neuroscience, 7(8):663–678.

Mehta, M., Lee, A., and Wilson, M. (2002). Role of experience and oscillations in transforming a rate code into a temporal code. Nature, 417(6890):741–746.

Mehta, M. R., Barnes, C. A., and McNaughton, B. L. (1997). Experience-dependent, asymmetric expansion of hippocampal place fields. Proceedings of the National Academy of Sciences, 94(16):8918–8921.

Mi, Y., Fung, C., Wong, K., and Wu, S. (2014). Spike frequency adaptation implements anticipative tracking in continuous attractor neural networks. Advances in neural information processing systems, 27.

Muessig, L., Lasek, M., Varsavsky, I., Cacucci, F., and Wills, T. J. (2019). Coordinated emergence of hippocampal replay and theta sequences during post-natal development. Current Biology, 29(5):834–840.

O’keefe, J. and Burgess, N. (2005). Dual phase and rate coding in hippocampal place cells: theoretical significance and relationship to entorhinal grid cells. Hippocampus, 15(7):853–866.

O’Keefe, J. and Recce, M. L. (1993). Phase relationship between hippocampal place units and the eeg theta rhythm. Hippocampus, 3(3):317–330.

Pfeiffer, B. E. (2020). The content of hippocampal “replay”. Hippocampus, 30(1):6–18.

Romani, S. and Tsodyks, M. (2015). Short-term plasticity based network model of place cells dynamics. Hippocampus, 25(1):94–105.

Samsonovich, A. and McNaughton, B. L. (1997). Path integration and cognitive mapping in a continuous attractor neural network model. Journal of Neuroscience, 17(15):5900–5920.

Schmidt, R., Diba, K., Leibold, C., Schmitz, D., Buzsáki, G., and Kempter, R. (2009). Single-trial phase precession in the hippocampus. Journal of Neuroscience, 29(42):13232–13241.

Skaggs, W. E., McNaughton, B. L., Wilson, M. A., and Barnes, C. A. (1996). Theta phase precession in hippocampal neuronal populations and the compression of temporal sequences. Hippocampus, 6(2):149–172.

Stewart, M. and Fox, S. E. (1990). Do septal neurons pace the hippocampal theta rhythm? Trends in neurosciences, 13(5):163–169.

Treves, A. (2004). Computational constraints between retrieving the past and predicting the future, and the ca3-ca1 differentiation. Hippocampus, 14(5):539–556.

Tsodyks, M. (1999). Attractor neural network models of spatial maps in hippocampus. Hippocampus, 9(4):481– 489.

Tsodyks, M. and Sejnowski, T. (1995). Associative memory and hippocampal place cells. International journal of neural systems, 6:81–86.

Tsodyks, M. V., Skaggs, W. E., Sejnowski, T. J., and McNaughton, B. L. (1996). Population dynamics and theta rhythm phase precession of hippocampal place cell firing: a spiking neuron model. Hippocampus, 6(3):271– 280.

Van Der Meer, M. A. and Redish, A. D. (2011). Theta phase precession in rat ventral striatum links place and reward information. Journal of neuroscience, 31(8):2843–2854.

Van Strien, N., Cappaert, N., and Witter, M. (2009). The anatomy of memory: an interactive overview of the parahippocampal–hippocampal network. Nature reviews neuroscience, 10(4):272–282.

Wang, M., Foster, D. J., and Pfeiffer, B. E. (2020). Alternating sequences of future and past behavior encoded within hippocampal theta oscillations. Science, 370(6513):247–250.

Wang, X.-J. (2002). Pacemaker neurons for the theta rhythm and their synchronization in the septohippocampal reciprocal loop. Journal of neurophysiology, 87(2):889–900.

Wikenheiser, A. M. and Redish, A. D. (2015). Hippocampal theta sequences reflect current goals. Nature neuroscience, 18(2):289–294.

Wu, S., Hamaguchi, K., and Amari, S.-i. (2008). Dynamics and computation of continuous attractors. Neural computation, 20(4):994–1025.

Wu, S., Wong, K. M., Fung, C. A., Mi, Y., and Zhang, W. (2016). Continuous attractor neural networks: candidate of a canonical model for neural information representation. F1000Research, 5.

Yamaguchi, Y. (2003). A theory of hippocampal memory based on theta phase precession. Biological cybernetics, 89(1):1–9.

Yamaguchi, Y., Aota, Y., McNaughton, B. L., and Lipa, P. (2002). Bimodality of theta phase precession in hippocampal place cells in freely running rats. Journal of neurophysiology, 87(6):2629–2642.

Zugaro, M. B., Monconduit, L., and Buzsáki, G. (2005). Spike phase precession persists after transient intrahippocampal perturbation. Nature neuroscience, 8(1):67–71.

