## Supplemental information text for "Firing rate adaptation affords place cell theta sweeps, phase precession and procession"

### Supplementary Materials for
### Firing rate adaptation in continuous attractor neural networks accounts for theta phase shift of hippocampal place cells

Tianhao Chu*, Zilong Ji*, Junfeng Zuo,Yuanyuan Mi, Wen-hao Zhang, Tiejun Huang, Daniel Bush, Neil Burgess, Si Wu†,

#### 1 The network model

We consider a continuous attractor neural network (CANN), in which neurons are uniformly distributed in an one-dimensional environment, mimicking place cells rearranged according to the locations of their firing fields on the linear track. Neurons in the CANN are connected with each other recurrently. Denote $U(x,t)$ the synaptic input received by neurons at location $x$, with $x \in (-\infty, \infty)$, and $r(x,t)$ the corresponding firing rate. The dynamics of the network are written as,

$$\tau \frac{dU(x,t)}{dt} = -U(x,t) + \rho \int_{-\infty}^{\infty} J(x,x') r(x',t)\, dx' - V(x,t) + I^{ext}(x,t), \tag{1}$$

$$\tau_v \frac{dV(x,t)}{dt} = -V(x,t) + mU(x,t), \tag{2}$$

$$r(x,t) = \frac{U(x,t)^2}{1 + k\rho \int_{-\infty}^{\infty} U(x',t)\, dx'}, \tag{3}$$

where $\tau$ is the time constant of $U(x,t)$, $\rho$ the neuron density and $I^{ext}(x,t)$ the external input. $J(x,x')$ represents the connection strength between neurons at $x$ and $x'$. $V(x,t)$ representing the effect of firing rate adaptation on this neuron. $\tau_v$ is the time constant of $V(x,t)$, and $m$ controls the adaptation strength. The parameter $k$ controls the amount of divisive normalization, reflecting the contribution of inhibitory neurons (not explicitly modeled) [1].

The connection profile between two neurons with firing fields at location $x$ and $x'$ is set as,

$$J(x,x') = \frac{J_0}{2\pi a} \exp\left[-\frac{(x-x')^2}{2a^2}\right], \tag{4}$$

where the parameter $J_0$ controls the amount of recurrent connection strength and $a$ represents the range of neuronal interactions. Notably, the recurrent connection between place cells is translation-invariant, that is, $J(x,x')$ is a function of $(x-x')$. This feature is crucial for the neutral stability of continuous attractor networks.

It is known that when there is no external input or firing rate adaptation ($m = 0$, $I^{ext} = 0$), the CANN can hold a continuous family of Gaussian-shaped stationary states [2, 3], called bump activities, as long as $k$ is smaller than a critical value $k_c = \rho J_0^2/(8\sqrt{2\pi}a)$. These bump states are expressed as $\overline{U}(x) = A_U \exp\left[-(x-z)^2/(4a^2)\right]$, with $z$ a free parameter representing the bump center and $A_U$ a constant representing the bump amplitude.

In general, although the state of the network is affected by external inputs and adaptation, the network bump can still be well approximated with Gaussian-like profiles [4]. Therefore, we assume the state of the network to be of the following Gaussain form,

$$\overline{U}(x,t) = A_u \exp\left\{-\frac{[x-z(t)]^2}{4a^2}\right\}, \tag{5}$$

$$\overline{V}(x,t) = A_v \exp\left\{-\frac{[x-z(t)+d(t)]^2}{4a^2}\right\}, \tag{6}$$

$$\overline{r}(x,t) = A_r \exp\left\{-\frac{[x-z(t)]^2}{2a^2}\right\}, \tag{7}$$

where $A_u$, $A_v$ and $A_r$ represent the amplitude of these Gaussian bumps. $z(t)$ is the center of $U(x,t)$ and $r(x,t)$. $d(t)$ denotes the distance between $U(x,t)$ and $V(x,t)$, and $d(t) > 0$ always holds, as the firing rate adaptation is a much slower dynamics lagging behind the neural dynamics. Here, we assume that the bump heights, i.e. $A_u$, $A_v$ and $A_r$, are all constants during the evolution of the neural dynamics.

To solve the specific solution of the network state, we need to substitute the general solution given by Eqs. 5-7 into the network dynamics Eqs. 1-3, we obtain

$$A_r = \frac{A_u^2}{1+k\rho\sqrt{2\pi}aA_u^2}, \tag{8}$$

$$\tau\left[A_u\frac{x-z}{2a^2}\frac{dz}{dt}+\frac{dA_u}{dt}\right]\mathcal{N}(z,2a) = (-A_u+\frac{\rho J_0}{\sqrt{2}}A_r)\mathcal{N}(z,2a) -A_v\mathcal{N}(z-d,2a)+I^{ext}(x,t), \tag{9}$$

$$\tau_v\left[A_v\frac{x-z+d}{2a^2}\frac{d(z-d)}{dt}+\frac{dA_v}{dt}\right]\mathcal{N}(z-d,2a) = -A_v\mathcal{N}(z-d,2a) +mA_u\mathcal{N}(z,2a), \tag{10}$$

where $\mathcal{N}(z,2a) = exp\left\{-[x-z]^2/4a^2\right\}$. Given $I^{ext}(x,t) = 0$, we can solve Eqs. 8-10 to get the solution of the network state. An important property of a CANN is that its dynamics is dominated by a few motion modes, as a consequence of the translation-invariant connections between neurons. We can therefore simplify the network dynamics significantly by projecting the network dynamics onto its dominating motion modes [3] (by projecting a function $f(x)$ onto a mode $u_n(x)$, it means to compute $\int_x f(x)u_n(x)dx$). Typically, projecting onto the first two motion modes is adequate (see next section).

For the bump $U(x,t)$, the first two motion modes are,

$$u_0(x,t) = \exp\left\{-\frac{[x-z(t)]^2}{4a^2}\right\}, \tag{11}$$

$$u_1(x,t) = [x-z(t)]\exp\left\{-\frac{[x-z(t)]^2}{4a^2}\right\}. \tag{12}$$

For the bump $V(x,t)$, the first two motion modes are,

$$v_0(x,t) = \exp\left\{-\frac{[x-z(t)+d(t)]^2}{4a^2}\right\}, \tag{13}$$

$$v_1(x,t) = [x-z(t)+d(t)]\exp\left\{-\frac{[x-z(t)+d(t)]^2}{4a^2}\right\}. \tag{14}$$

### 2 Deriving the network state when the external input does not exist ($I^{ext} = 0$)

#### 2.1 Static bump state of the network

We first analyze the condition for the network holding a static bump as its stationary state. In this case, the positions of bumps $U$ and $V$ remain unchanged, i.e., $dz/dt = 0$, and the discrepancy between them is zero, i.e., $d = 0$. Thus, Eqs. 9-10 can be simplified as,

$$\tau \frac{dA_u}{dt} = -A_u + \frac{\rho J_0}{\sqrt{2}} A_r - A_v, \tag{15}$$

$$\tau_v \frac{dA_v}{dt} = -A_v + mA_u. \tag{16}$$

Combining them with Eq. 8, we obtain the solution of the steady state of the network as

$$A_v = mA_u, \tag{17}$$

$$A_r = \frac{\sqrt{2}(1+m)}{\rho J_0} A_u, \tag{18}$$

$$A_u = \frac{\rho J_0 + \sqrt{\rho^2 J_0^2 - 8\sqrt{2\pi}(1+m)^2 k\rho a}}{4\sqrt{\pi}(1+m)k\rho a}. \tag{19}$$

To analyze the stability of this solution, we calculate the Jacobian matrix at this state, which is given by,

$$\mathbf{M} = \begin{pmatrix} \frac{1}{\tau}\left(-1 + \frac{\sqrt{2}\rho J_0 A_u}{(1+\sqrt{2\pi}k\rho a A_u^2)^2}\right) & -\frac{1}{\tau} \\ \frac{m}{\tau_v} & -\frac{1}{\tau_v} \end{pmatrix} \tag{20}$$

Denote the eigenvalues of the Jacobian matrix as $\lambda_1$ and $\lambda_2$. Therefore, the condition for the solution to be stable is that both eigenvalues are negative, which gives

$$\lambda_1 + \lambda_2 = \frac{1}{2}\left[-1 + \frac{\sqrt{2}A_u J_0 \rho}{(1+\sqrt{2\pi}k\rho a A_u^2)^2} - \frac{\tau}{\tau_v}\right] < 0, \tag{21}$$

$$\lambda_1 \lambda_2 = \frac{\tau}{\tau_v}\left(m + 1 - \frac{\sqrt{2}A_u J_0 \rho}{(1+\sqrt{2\pi}k\rho a A_u^2)^2}\right) > 0. \tag{22}$$

The above inequalities are satisfied when,

$$0 < k < k_{c1} = \frac{\rho J_0^2 (1+\frac{\tau}{\tau_v})(1+2m-\frac{\tau}{\tau_v})}{8\sqrt{2\pi}a(1+m)^4}, \tag{23}$$

$$0 < k < k_{c2} = \frac{\rho J_0^2}{8\sqrt{2\pi}a(1+m)^2}. \tag{24}$$

It is easy to check that $k_{c2} < k_{c1}$, so the condition for the network to hold static bumps as its steady state is $0 < k < k_{c2}$.

#### 2.2 Travelling wave state of the network

We further analyze the condition for the network holding a continuously moving bump (travelling wave) as its steady state. In this state, the bump moves at a constant speed, and the center position is expressed as,

$$z(t) = v_{int} t, \tag{25}$$

where $v_{int}$ is called the intrinsic speed of the bump activity. Since the bump height is roughly unchanged and the discrepancy $d$ is a constant, Eqs. 9-10 can be simplified as,

$$\tau\left(A_u \frac{x-z}{2a^2} v_{int}\right)\mathcal{N}(z,2a) = (-A_u + \frac{\rho J_0}{\sqrt{2}}A_r)\mathcal{N}(z,2a) - A_v \mathcal{N}(z-d,2a), \tag{26}$$

$$\tau_v\left(A_v \frac{x-z+d}{2a^2} v_{int}\right)\mathcal{N}(z-d,2a) = -A_v \mathcal{N}(z-d,2a) + mA_u \mathcal{N}(z,2a). \tag{27}$$

In order to obtain the solution of variables of $A_u, A_v, A_r, d$ and $v_{int}$, we reduce the dimensionality of the network dynamics by projecting Eqs. 26-27 onto the dominant motion modes $u_0(x), u_1(x), v_0(x), v_1(x)$. First, projecting both sides of Eq. 26 onto the motion mode $u_0(x)$ (expressed in Eq.11), we obtain

$$Left-side = 0,$$

$$Right-side = (-A_u + \frac{\rho J_0}{\sqrt{2}}A_r)\sqrt{2\pi}a - A_v \exp(-\frac{d^2}{8a^2})\sqrt{2\pi}a.$$

Equating both sides, we have

$$-A_u + \frac{\rho J_0}{\sqrt{2}} A_r - A_v \exp(-\frac{d^2}{8a^2}) = 0. \tag{28}$$

Similarly, projecting Eq. 26 onto the motion mode $u_1(x)$ (expressed in Eq.12) and equating both sides, we obtain

$$\tau A_u v_{int} = d A_v \exp(-\frac{d^2}{8a^2}). \tag{29}$$

Again, projecting both sides of Eq. 27 onto the motion modes $u_0(x)$ and $u_1(x)$, respectively, and equating both sides, we obtain

$$\frac{d}{4a^2}\tau_v A_v \exp(-\frac{d^2}{8a^2}) v_{int} = -A_v \exp(-\frac{d^2}{8a^2}) + m A_u, \tag{30}$$

$$\tau_v (1 - \frac{d^2}{4a^2}) v_{int} = d. \tag{31}$$

Combining Eq.8 and Eqs.28-31, we obtain the values of $A_u, A_v, A_r, d$ and $v_{int}$ in the travelling wave state as follows

$$A_u = \frac{\rho J_0 + \sqrt{\rho^2 J_0^2 - 8\sqrt{2\pi} k\rho a (1 + \sqrt{\frac{m\tau}{\tau_v}})^2}}{4\sqrt{\pi} k\rho a (1 + \sqrt{\frac{m\tau}{\tau_v}})}, \tag{32}$$

$$A_v = \frac{\rho J_0 + \sqrt{\rho^2 J_0^2 - 8\sqrt{2\pi} k\rho a (1 + \sqrt{\frac{m\tau}{\tau_v}})^2}}{2\sqrt{2\pi} k\rho^2 a J_0}, \tag{33}$$

$$A_r = \sqrt{\frac{m\tau}{\tau_v}} \exp\left[\frac{1 - \sqrt{\frac{\tau}{m\tau_v}}}{2}\right] \frac{\rho J_0 + \sqrt{\rho^2 J_0^2 - 8\sqrt{2\pi} k\rho a (1 + \sqrt{\frac{m\tau}{\tau_v}})^2}}{4\sqrt{\pi} k\rho a (1 + \sqrt{\frac{m\tau}{\tau_v}})}, \tag{34}$$

$$d = 2a\sqrt{1 - \sqrt{\frac{\tau}{m\tau_v}}}, \tag{35}$$

$$v_{int} = \frac{2a}{\tau_v}\sqrt{\frac{m\tau_v}{\tau} - \sqrt{\frac{m\tau_v}{\tau}}}. \tag{36}$$

It is straightforward to check from Eq. 36 that, to obtain a traveling wave state, $v_{int}$ should be a real positive value. This give the condition that

$$m > \frac{\tau}{\tau_v}. \tag{37}$$

Eqs. 24&37 gives the phase diagram of the network state when external input does not exist, as shown in Fig.2g in the main text.

### 3 Deriving the oscillatory tracking state of the network when the external input is applied ($I^{ext} \neq 0$)

The external input is given by

$$I^{ext} = \alpha \exp\left[-\frac{(x - v_{ext}t)^2}{4a^2}\right], \tag{38}$$

where $\alpha$ is the input strength and $v_{ext}$ is the speed of the external input, mimicking the moving speed of the artificial animal.

The network state is mainly affected by two competing factors: one is the intrinsic mobility of the network originated from the firing rate adaptation, which drives the bump to move spontaneously (see SI.Sec. 2.2 above); the other is the extrinsic mobility driven by the external input, which drives the bump to move at the same speed of $v_{ext}$. The competition between these two factors leads to three tracking states of the network: travelling wave, oscillatory tracking, and smooth tracking. Among the three states, the travelling wave state and the smooth tracking state are similar to the two cases we described in the previous section. Here we only focus on the analytical derivation of the oscillatory tracking state. By numerically simulating the attractor network, we find that the bump center position can be roughly expressed as a sinusoidal moving wave give by,

$$z(t) = c_0 sin(\omega t) + d_0 + v_{ext} t, \tag{39}$$

where $c_0$ and $\omega$ represent the amplitude and frequency of the sinusoidal wave, respectively, and $d_0$ denotes the offset between the center of the activity bump and the center of the external input. Substituting the form of external input expressed in Eq. 38 into Eqs. 9-10, we obtain

$$\begin{aligned} \tau\left(A_u \frac{x-z}{2a^2}\frac{dz}{dt}\right)\mathcal{N}(z,2a) &= (-A_u + \frac{\rho J_0}{\sqrt{2}}A_r)\mathcal{N}(z,2a) - A_v\mathcal{N}(z-d,2a) \\ &\quad +\alpha\mathcal{N}(v_{ext}t,2a), \qquad (40) \\ \tau_v\left(A_v \frac{x-z+d}{2a^2}\frac{d(z-d)}{dt}\right)\mathcal{N}(z-d,2a) &= -A_v\mathcal{N}(z-d,2a) + mA_u\mathcal{N}(z,2a). \qquad (41) \end{aligned}$$

Again, to get the solution of variables of $A_u, A_v, A_r, c_0, \omega, d_0$ we reduce the dimensionality of the network dynamics by projecting Eqs. 40-41 onto the dominant motion modes $u_0(x), u_1(x), v_0(x), v_1(x)$. Projecting Eq. 40 onto $u_0$ and $u_1$, respectively, gives

$$\begin{aligned} -A_u + \frac{\rho J_0}{\sqrt{2}}A_r + \alpha\exp(-\frac{s^2}{8a^2}) &= A_v\exp(-\frac{d^2}{8a^2}), \qquad (42) \\ dA_v\exp(-\frac{d^2}{8a^2}) - \alpha s\exp(-\frac{s^2}{8a^2}) &= \tau A_u\frac{dz}{dt}, \qquad (43) \end{aligned}$$

where $s(t) = c_0\sin(\omega t) + d_0$ denotes the offset between $U(x,t)$ and $I_{ext}(x,t)$. For clearance, we denote $A_{temp} = A_v\exp(-d^2/8a^2)$ hereafter.

We first solve the dynamics of the distance $d(t)$, i.e., the distance between $U(x,t)$ and $V(x,t)$. To do this, we substitute Eqs. 39&42 into Eq. 43 and obtain

$$d(t) = \frac{1}{A_{temp}}\left[\tau A_u(v + c_0\omega\cos\omega t) + \alpha s\exp(-\frac{s^2}{8a^2})\right]. \tag{44}$$

Since $s \ll 2a$ generally holds, we have $\exp(-s^2/8a^2) \approx 1$. With this approximation, the equation above can be re-written as

$$d(t) = A_0\sin(\omega t + \beta) + B_0, \tag{45}$$

where the parameters are solved as,

$$\begin{aligned} \beta &= \arccos\left(\frac{\alpha}{\sqrt{\tau^2 A_u^2\omega^2 + \alpha^2}}\right), \qquad (46) \\ A_0 &= \frac{c_0\sqrt{\tau^2 A_u^2\omega^2 + \alpha^2}}{A_{temp}}, \qquad (47) \\ B_0 &= \frac{\tau A_u v + \alpha d_0}{A_{temp}}. \qquad (48) \end{aligned}$$

Again, projecting Eq. 41 onto $v_0$ and $v_1$, respectively, gives

$$\begin{aligned} A_v &= mA_u\exp(-\frac{d^2}{8a^2}), \qquad (49) \\ \tau_v A_v\left[\frac{\mathrm{d}z(t)}{\mathrm{d}t} - \frac{\mathrm{d}d(t)}{\mathrm{d}t}\right] &= mA_u\exp(-\frac{d^2}{8a^2})d(t). \qquad (50) \end{aligned}$$

To solve the expression of $A_u$, we substitute Eqs. 8&49 into Eq. 42 and obtain

$$-A_u + \frac{\rho J_0}{\sqrt{2}}\frac{A_u^2}{1+\sqrt{2\pi}ak\rho A_u^2} + \alpha\exp(-\frac{s^2}{8a^2}) = mA_u\exp(-\frac{d^2}{4a^2}). \tag{51}$$

Since $s \ll 2a$ and $d \ll 2a$, the approximations of $\exp(-s^2/8a^2) \approx 1$ and $\exp(-d^2/4a^2) \approx 1$ hold, and the above equation can be simplified as,

$$(m+1)A_u - \frac{\rho J_0}{\sqrt{2}} \frac{A_u^2}{1+\sqrt{2\pi}ak\rho A_u^2} - \alpha = 0. \tag{52}$$

We can rearrange Eq. 52 into a general cubic equation of $A_u$, which is written as,

$$a_3 A_u^3 + a_2 A_u^2 + a_1 A_u + a_0 = 0, \tag{53}$$
$$a_3 = \sqrt{2\pi}(m+1)ak\rho, \tag{54}$$
$$a_2 = -\sqrt{2\pi}ak\rho\alpha - \frac{\rho J_0}{\sqrt{2}}, \tag{55}$$
$$a_1 = m+1, \tag{56}$$
$$a_0 = -\alpha, \tag{57}$$

It's easy to check that Eq. 53 only have one real solution, which is

$$A_u = \left[-\frac{q}{2} + \sqrt{(\frac{q}{2})^2 + (\frac{p}{3})^3}\right]^{1/3} + \left[-\frac{q}{2} - \sqrt{(\frac{q}{2})^2 + (\frac{p}{3})^3}\right]^{1/3}, \tag{58}$$
$$q = \frac{3a_3a_1 - a_2^2}{3a_3^2}, \tag{59}$$
$$p = \frac{27a_3^2a_0 - 9a_3a_2a_1 + 2a_2^3}{27a_3^3}. \tag{60}$$

The analytical solution of $A_u$ given by Eqs. 58-60 is very complicated, However, by numerically simulating the network, we find that $\sqrt{2\pi}ak\rho A_u^2 \gg 1$ can be hold when the network is at the oscillatory tracking state. This gives $A_u^2/(1+\sqrt{2\pi}ak\rho A_u^2) \approx 1/\sqrt{2\pi}ak\rho$. Therefore, we can simplify the expression of $A_u$ to

$$A_u = \frac{J_0 + 2\sqrt{\pi}ak\alpha}{2\sqrt{\pi}ak(1+m)}. \tag{61}$$

To solve $\omega, d_0$ and $c_0$, we further substitute Eq. 49 into Eq. 50 and obtain

$$\tau_v \frac{\mathrm{d}z(t)}{\mathrm{d}t} = d(t) + \tau_v \frac{\mathrm{d}d(t)}{\mathrm{d}t}. \tag{62}$$

The above equation can be expanded by substituting Eqs. 39 & 45 into Eq. 62 which gives

$$\tau_v(v + c_0\omega\cos\omega t) = A_0\left[\sin(\omega t+\beta) + \omega\tau_v\cos(\omega t+\beta)\right] + B_0.$$

Using the trigonometric transformation formula, we can rewrite the above equation as

$$\tau_v v + \tau_v c_0\omega\sin(\omega t + \frac{\pi}{2}) = A_0\sqrt{1+\omega^2\tau_v^2}\sin(\omega t+\beta+\gamma) + B_0, \tag{63}$$

where $\gamma$ is given by

$$\gamma = \arccos(\frac{1}{\sqrt{\tau_v^2\omega^2}}). \tag{64}$$

Equating two sides of Eq. 63, we have,

$$\tau_v v = B_0, \tag{65}$$
$$\frac{\pi}{2} = \beta + \gamma, \tag{66}$$
$$\tau_v c_0\omega = A_0\sqrt{1+\omega^2\tau_v^2}. \tag{67}$$

Now we combine the three equations given by Eqs. 65-67 to get the solutions of $c_0, \omega$ and $d_0$. Substituting Eq. 48 into Eq. 65, we obtain

$$d_0 = \frac{\tau_v v A_{temp} - \tau v A_u}{\alpha}. \tag{68}$$

Applying the cosine function to both sides of Eq. 66, we obtain

$$\cos(\beta+\gamma) = \cos\gamma\cos\beta - \sin\gamma\sin\beta = 0. \tag{69}$$

Substituting Eqs. 46 & 64 into 69, we have

$$\omega^2 = \frac{\alpha}{\tau\tau_v A_u}. \tag{70}$$

Combining Eq. 70 with Eq. 61, we obtain the expression for the oscillating frequency $\omega$, that is,

$$\omega = \sqrt{\frac{2\sqrt{\pi}\alpha ak(1+m)}{\tau\tau_v(J_0 + 2\sqrt{\pi}ak\alpha)}}. \tag{71}$$

Substituting Eqs. 47 & 70 into Eq. 67, and taking square for both sides, we have

$$\frac{(\tau^2 A_u^2\omega^2 + \alpha^2)}{A_{temp}^2}(1+\omega^2\tau_v^2) = \tau_v^2\omega^2. \tag{72}$$

Solving the above equation for $A_{temp}$, we get

$$A_{temp} = \frac{\tau A_u + \alpha\tau_v}{\tau_v}. \tag{73}$$

Substituting Eq. 73 into Eq. 68, we can get the expression for the average offset $d_0$ between the bump center and the external input center, which is

$$d_0 = \tau_v v. \tag{74}$$

Since $A_{temp} = mA_u \exp\left[-d(t)^2/(4a^2)\right]$ varies across time, we take the approximation

$$A_{temp} = mA_u \exp\left[-\overline{d(t)^2}/(4a^2)\right], \tag{75}$$

with $\overline{d(t)^2}$ the time-averaged value, which is calculated to be,

$$\overline{d(t)^2} = \frac{1}{T}\int_0^T d^2(t)dt = \frac{\alpha\tau_v}{2(\tau A_u + \alpha\tau_v)}c_0^2 + \tau_v^2 v^2. \tag{76}$$

Substituting Eqs. 73 & 76 into Eq. 75, we obtain the expression for the oscillating amplitude $c_0$, that is,

$$c_0 = \sqrt{\frac{2(\tau A_u + \alpha\tau_v)}{\alpha\tau_v}\left[4a^2(\ln\frac{\tau_v mA_u}{\tau A_u + \alpha\tau_v}) - \tau_v^2 v^2\right]}, \tag{77}$$

Substituting Eq. 73 into Eq. 49 and utilizing the condition of $\exp(-d^2/8a^2) = \sqrt{A_{temp}/mA_u}$, we obtain the expression for the bump height of $V(x,t)$

$$A_v = \sqrt{\left(\frac{\tau A_u + \alpha\tau_v}{\tau_v}\right)mA_u}. \tag{78}$$

Overall, combining Eqs.8, 61, 74, 71, 77,& 78, we get the solutions of all variables in the oscillatory tracking state, which are expressed as

$$A_u = \frac{J_0 + 2\sqrt{\pi}ak\alpha}{2\sqrt{\pi}ak(1+m)}, \tag{79}$$

$$A_r = \frac{A_u^2}{1+\sqrt{2\pi}ak\rho A_u^2}, \tag{80}$$

$$A_v = \sqrt{\left(\frac{\tau A_u + \alpha\tau_v}{\tau_v}\right)mA_u}, \tag{81}$$

$$c_0 = A_v\sqrt{\frac{2}{\alpha mA_u}\left[8a^2\ln\frac{mA_u}{A_v} - \tau_v^2 v^2\right]}, \tag{82}$$

$$d_0 = \tau_v v, \tag{83}$$

$$\omega = \sqrt{\frac{\alpha}{\tau\tau_v A_u}}. \tag{84}$$

It is noteworthy that the theoretical solutions given by Eqs. 82-84 exist only if the sweep amplitude $c_0$ given by Eq. 82 is of real value. This gives the condition for the CANN to be in the oscillatory tracking state, which is

$$8a^2 \ln \frac{mA_u}{A_v} > \tau_v^2 v^2. \tag{85}$$

We carry out numerical simulations to verify our theoretical results, including the theoretical solutions of the mean offset $d_0$ given by Eq. 83 (see SI Fig.2a-c) and the theoretical boundary that separates the smooth tracking state and the oscillatory tracking state given by Eq. 85 (see SI Fig. 2d). The results show that our theoretical analysis agrees well with the simulation results.

### 4 Oscillatory tracking in the 2-dimensional CANN – modeling theta sweeps in the open field environment

#### 4.1 Model description

In the two-dimensional CANN, neurons are uniformly distributed on a rectangular neuronal sheet arranged according to the locations of their firing fields. Denote $U(\mathbf{x}, t)$ as the synaptic input to the neuron at location $\mathbf{x}$, with $\mathbf{x} = (x_1, x_2)$ and $x_1, x_2 \in (-\infty, \infty)$, and $r(\mathbf{x}, t)$ as the corresponding firing rate. The dynamics of $U(\mathbf{x}, t)$ is determined by its own relaxation, the recurrent inputs from other neurons, and the firing rate adaptation, which is written as,

$$\tau \frac{\partial U(\mathbf{x}, t)}{\partial t} = -U(\mathbf{x}, t) + \rho \int_{\mathbf{x}'} J(\mathbf{x}, \mathbf{x}') r(\mathbf{x}', t) \, \mathrm{d}\mathbf{x}' - V(\mathbf{x}, t) + \sigma_U \xi_U(\mathbf{x}, t). \tag{86}$$

Here, $\tau$ is the time constant of synaptic current and $\rho$ is the neuronal density. The recurrent connection is defined as$J(\mathbf{x}, \mathbf{x}') = J_0/(2\pi a^2) \exp\left[-\|\mathbf{x} - \mathbf{x}'\|^2/(2a^2)\right]$, with $\|\mathbf{x} - \mathbf{x}'\|^2 = (x_1 - x_1')^2 + (x_2 - x_2')^2$ which is translation-invariant on the neuronal sheet (SIFig. 6a). The nonlinear relationship between the firing rate $r(\mathbf{x}, t)$ and the synaptic input $U(\mathbf{x}, t)$ is implemented by the divisive normalization, which is

$$r(\mathbf{x}, t) = \frac{U^2(\mathbf{x}, t)}{1 + k\rho \int_{\mathbf{x}'} U^2(\mathbf{x}', t) \, \mathrm{d}\mathbf{x}'}, \tag{87}$$

where $k$ controls the normalization strength. In the neural system, divisive normalization could be implemented by shunting inhibition [1]. The term $V(\mathbf{x}, t)$ on the right-hand side of Eq. (86) represents the effect of firing rate adaptation, with the dynamics written as

$$\tau_v \frac{\partial V(\mathbf{x}, t)}{\partial t} = -V(\mathbf{x}, t) + mU(\mathbf{x}, t), \tag{88}$$

where $\tau_v$ is the time constant, and $m$ is the adaptation strength.

#### 4.2 Theta sweeps in the 2-dimensional CANN

We study the network dynamics when an moving external input is applied to the network, which is written as,

$$I^{ext} = \alpha \exp\left[-\frac{||\mathbf{x} - \mathbf{v_{ext}} t||^2}{4a^2}\right], \tag{89}$$

where $\mathbf{v_{ext}} = (v_x, v_y)$ represents the speed of the external input and $\alpha$ represents the input strength. We consider one simple case, where the external input is moving on a straight line along the X-axis, i.e., $v_y = 0$. We find that, similar to 1D CANN, when the input strength $\alpha$ and adaptation strength $m$ is set appropriately, the bump activity also oscillates around the moving input along the moving direction, which give rise to the alternative forward and reverse theta sequences along moving direction of the external input (SIFig. 6b). The heatmap of the phase shift of the probe neuron which is located at $x = 0, y = 0$ is shown in SIFig. 6c.

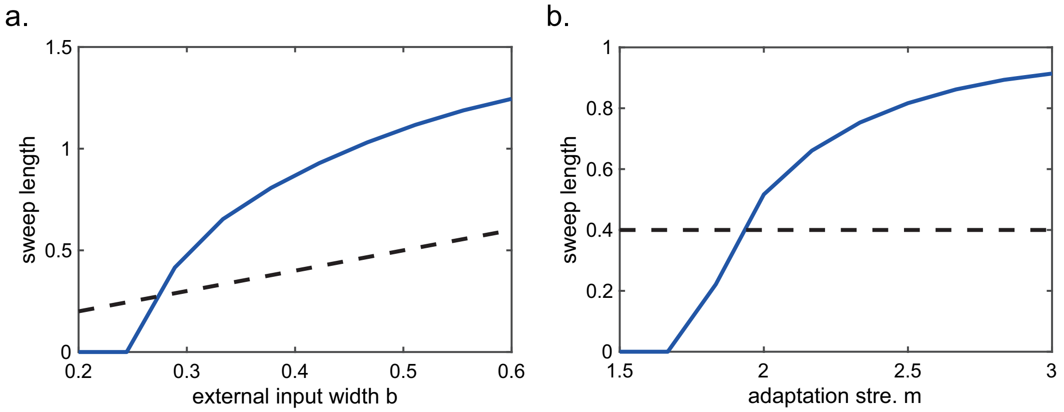


Figure 1: Sweep length is not bounded by the external input width. **a**, the sweep length is positively but not linearly related with the external input width. **b**, with fixed external input width, increasing the adaptation strength the sweep length can exceed the external input width. This figure relates to Fig. 2 in the main text.

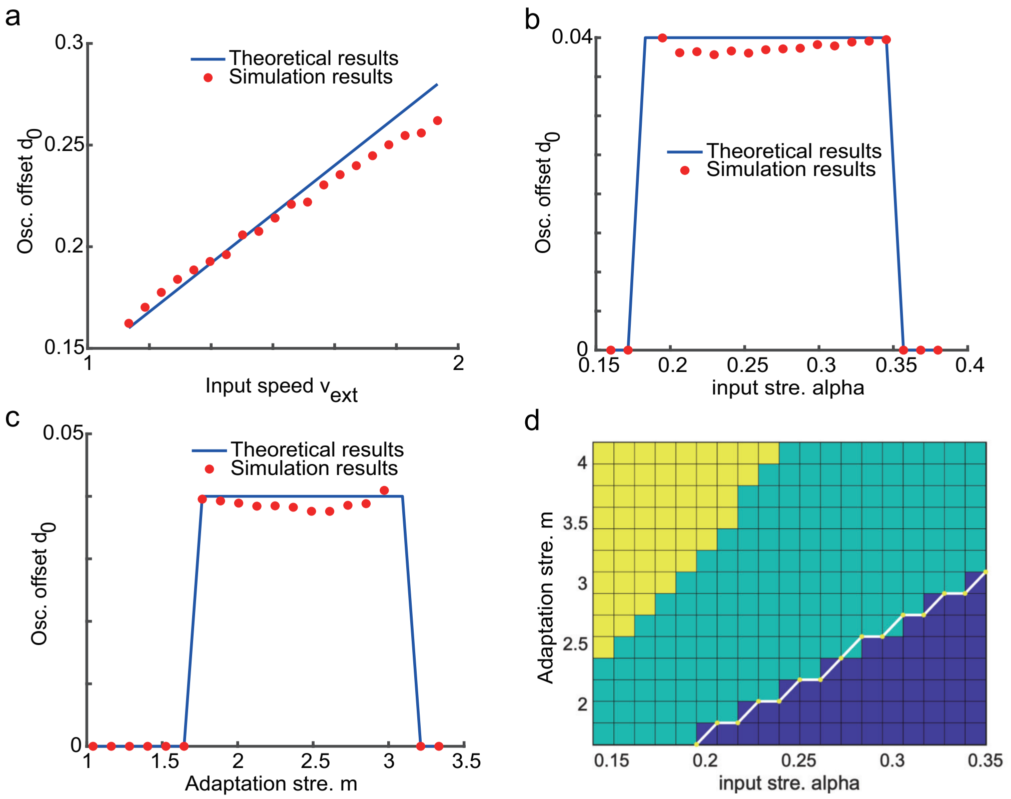


Figure 2: Verifying theoretical results with numerical simulations. **a-c**, Simulation results of the average offset $d_0$ as a function of $v_{ext}$, $\alpha$ and $m$, respectively. **d**, The phase diagram of network states. The yellow area represents the traveling wave state, the green area represents the oscillatory tracking state and the blue area represents the smooth tracking state. The white line represents the theoretical boundary given by Eq. 85. The parameters used in simulations are: $k = 5$, $J_0 = 1$, $a = 0.4$, $N = 512$, $\tau = 3$ ms, $\tau_v = 144$ ms, $\rho = 20.37$. This figure relates to Fig. 2 in the main text.

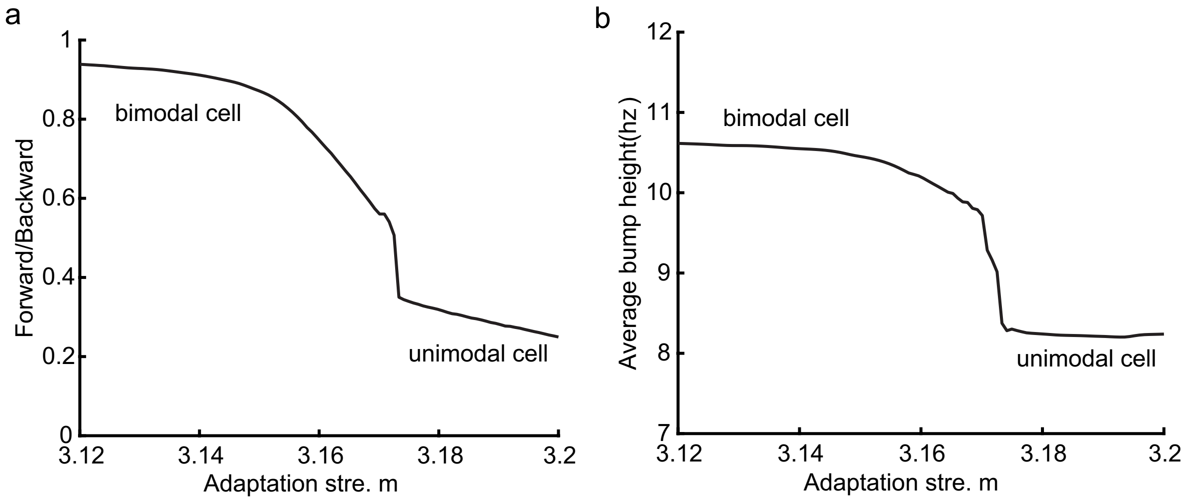


Figure 3: Activity bump height as a function of the adaptation strength. **a**, The ratio between the average bump height during forward window and the average bump height during backward window as function of the adaptation $m$. When the adaptation strength is relatively small, the mean firing rate of place cells are approximately the same in the forward window as in the backward window. And the place cells exhibit bimodal cell properties. As the adaptation strength gets larger, the mean firing rates in the backward window gradually decrease and the place cells tend to exhibit firing properties more like unimodal cells. **b**, The average bump height as a function of the adaptation strength $m$. Our model predicts that the bimodal cells fire at higher frequency than unimodal cells which can be testable in future experiments. The parameters are:$\alpha = 0.19$, $k = 5$,$J_0 = 1$, $a = 0.4$, $N = 512$, $\tau = 3ms$, $\tau_v = 144ms$, $\rho = 20.37$, $v_{ext} = 0.51m/s$. This figure relates to Fig. 4 in the main text.

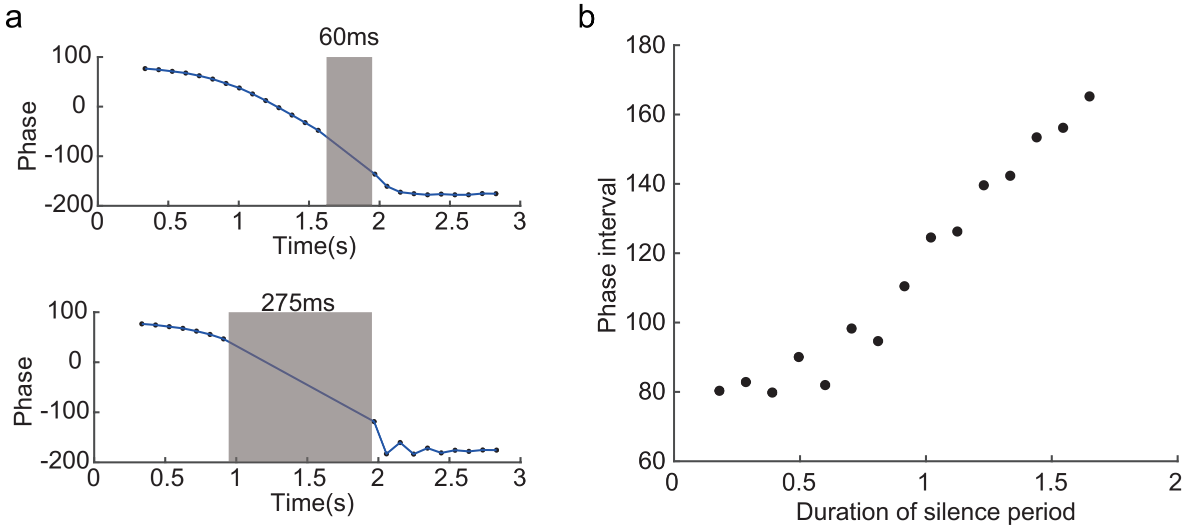


Figure 4: Persistent phase shift with variable silencing periods. **a**, Two examples of the persisting phase shift after transient silencing. Upper panel: the silencing duration is $60ms$. Upper panel: the silencing duration is $275ms$. **b**, The phase interval before and after the silencing as a function of the duration of the silencing. The phase interval gradually increases with the silencing duration. The parameters are: $\alpha = 0.19$,$m = 3.23$, $a = 0.4$,$k = 5$,$J_0 = 1$, $N = 512$, $\tau = 3ms$, $\tau_v = 144ms$, $\rho = 20.37$, $v_{ext} = 1m/s$. This figure relates to Fig. 6 in the main text.

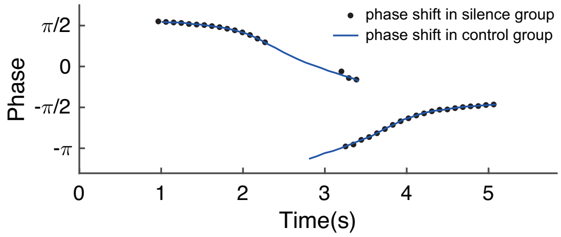


Figure 5: Persistent bimodal phase shift after transient silencing.The parameters are: $\alpha = 0.19$,$m = 3.03$, $a = 0.4$,$k = 5$,$J_0 = 1$, $N = 512$, $\tau = 3ms$, $\tau_v = 144ms$, $\rho = 20.37$, $v_{ext} = 0.3m/s$. This figure relates to Fig. 6 in the main text.

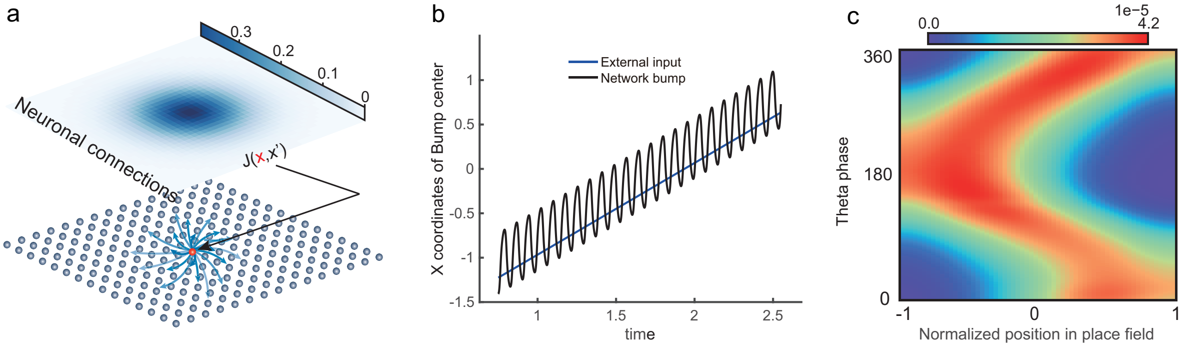


Figure 6: Theta sweeps and theta phase shift in a 2D CANN. **a**, An demonstration of the 2D CANN. **b**, The trajectory of the bump center and external input center when the input is moving along the X-axis in the 2D CANN. **c**, Theta phase as a function of the normalized position of the animal in place field, averaged over all place cells that are placed on the X-axis. $-1$ represents that the animal just entries the place field, and 1 represents that the animal is about to leave the place field. The parameters are: $\alpha = 0.2$,$m = 4$, $k = 6$,$J_0 = 1$, $a = 0.3$, $N = 16384$, $\tau = 3ms$, $\tau_v = 150ms$, $\rho = 415.01$, $v_{ext} = 1m/s$. This figure relates to Fig. 2 in the main text.

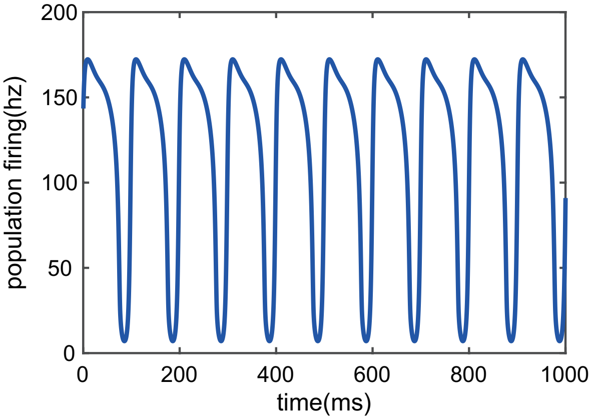


Figure 7: Theta oscillation of the population activities during the theta sweep state. This figure relates to Fig. 4 in the main text.

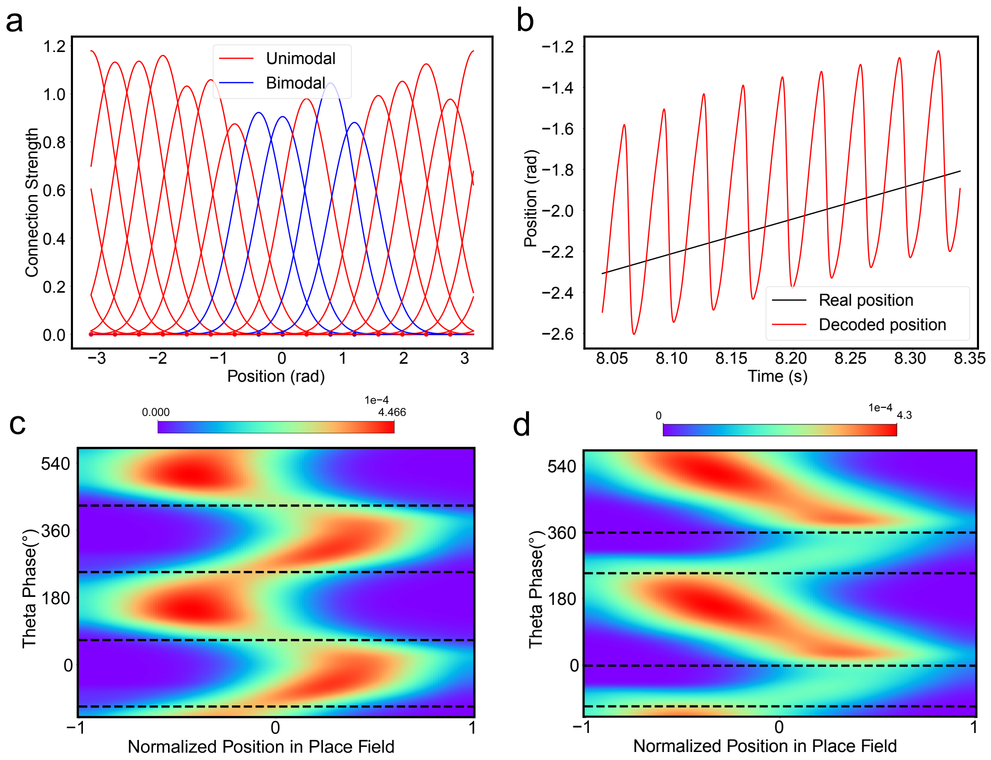


Figure 8: A-CANN with heterogeneous connection strength generate oscillatory tracking to account for theta phase shift. **a**, The synaptic connection strength profile of the neurons in the network. The blue lines represent the synaptic strengths of the neurons which turn out to be bimodal neurons, while the red lines represent the unimodal neurons. **b**, The oscillatory tracking trajectory of the bump center. **c**, The phase shift distribution of the bimodal cells. **d**, The phase shift distribution of the unimodal cells. The variations of the connection strength is 0.1, the average value is 1. Other parameters are the same with the figure 3 and 4 in the main text.

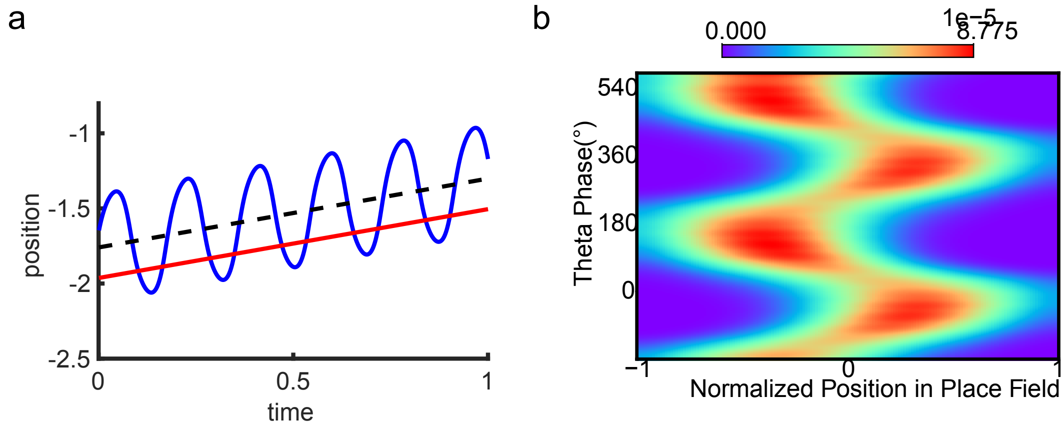


Figure 9: Oscillatory tracking behavior accounts for theta phase shift with $tau = 10ms$. **a**, oscillatory tracking behavior. **b**, bimodal phase shift of one example neuron. $\tau = 10ms., \tau_v = 480ms, \alpha = 0.3, m = 6$, Other parameters are set equal with the figure 2 in the main text. This figure relates to Fig. 2 in the main text.

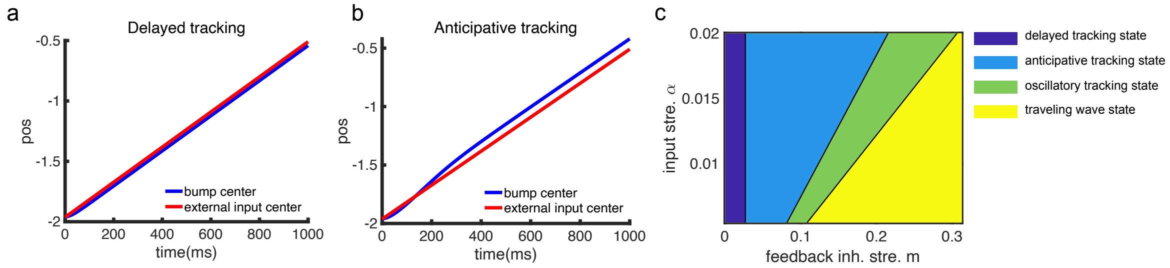


Figure 10: Spatio-temporal tracking dynamics when the adaptation strength is low (start from 0). **a**, the tracking behavior when the adaptation strength $m = 0$ ($\alpha = 0.02$). The network bump can generate a bump to track the external input stimuli smoothly but with a contant lagging distance which is proportional to the time constants $\tau$ and the external input speed $v_{ext}$. **b**, the tracking behavior when the adaptation strength $m = 0.1$ ($\alpha = 0.02$). Thanks to the intrinsic mobility introduced by SFA, the network bump anticipatively track the external input stimuli with a contant leading distance which is proportional to the adaptation strength $m$ and the external input speed $v_{ext}$. **c**, a phase diagram that summarizes the spatiotemporal patterns of the A-CANN tracking behavior. Other parameters are set equal with the figure 2 in the main text. This figure relates to Fig. 2 in the main text.

### References

[1] Simon J Mitchell and R Angus Silver. Shunting inhibition modulates neuronal gain during synaptic excitation. *Neuron*, 38(3):433–445, 2003.

[2] CC Alan Fung, KY Michael Wong, He Wang, and Si Wu. Dynamical synapses enhance neural information processing: gracefulness, accuracy, and mobility. *Neural computation*, 24(5):1147–1185, 2012.

[3] CC Alan Fung, KY Michael Wong, and Si Wu. A moving bump in a continuous manifold: a comprehensive study of the tracking dynamics of continuous attractor neural networks. *Neural Computation*, 22(3):752–792, 2010.

[4] Yuanyuan Mi, CC Fung, KY Wong, and Si Wu. Spike frequency adaptation implements anticipative tracking in continuous attractor neural networks. *Advances in neural information processing systems*, 27, 2014.
